# Ubiquitin-mediated stabilization of SlPsbS regulates low night temperature tolerance in tomatoes

**DOI:** 10.1101/2024.02.21.581502

**Authors:** Jiazhi Lu, Junchi Yu, Pengkun Liu, Jiamao Gu, Yu Chen, Tianyi Zhang, Jialong Li, Taotao Wang, Wenqiang Yang, Rongcheng Lin, Feng Wang, Mingfang Qi, Tianlai Li, Yufeng Liu

**Author notes:** Correspondence: Tianlai Li, Yufeng Liu.

## Abstract

Non-photochemical quenching (NPQ) plays a fundamental role in safely operating photosynthesis under low night temperatures (LNT). PsbS protein is essential for the rapid induction of NPQ, and its stability is often affected by adverse environmental conditions. However, the regulatory mechanism for the stability of PsbS or chloroplast proteins remains to be fully characterized. We showed that LNT decreased NPQ levels and SlPsbS protein abundance in tomato leaves. COP9 signalosome subunit 5A (SlCSN5A) facilitated SlPsbS ubiquitination and degradation in the cytosol. Further, tomato chloroplast vesiculation (SlCV) was activated by LNT. Under LNT, SlCV targeted the chloroplasts and induced the formation of CV-containing vesicles (CCVs) containing SlPsbS, which were exported from the chloroplasts. Subsequently, SlCV and SlPsbS contact SlCSN5A in the cytosol and are ubiquitinated and degraded. Genetic evidence demonstrated that overexpression of SlCV aggravated SlPsbS protein degradation, whereas silencing of SlCSN5 and SlCV delayed LNT-induced NPQ reduction and SlPsbS protein turnover. This study provides evidence that CSN5A is associated with chloroplast proteins, and reveals a ubiquitin-dependent degradation pathway of chloroplast proteins co-mediated by CV and CSN5A, thereby providing new insights into the regulation of chloroplast protein stability under stress conditions.

## INTRODUCTION

Low temperature is an environmental factor that severely decreases crop growth and productivity. Chloroplasts have been suggested to act as central sensors of and responders to stress and are the main site of oxidative damage in plant leaves under low temperatures. This may lead to photoinhibition of photosynthesis (Gao et al., 2022). Plants have evolved strategies to rapidly respond to stress, for instance, non-photochemical quenching (NPQ) is broadly considered a major factor in the rapid regulation of light harvesting to protect the photosystem II (PSII) reaction centers against photodamage (Ghosh et al., 2023). PsbS protein was identified as a major induced protein for NPQ activation in plants (Correa-Galvis et al., 2015). PsbS protein accumulates rapidly and transiently under stress to induce NPQ, while prolonged or severe stress accelerates its degradation (Nawrocki et al., 2020; Redekop et al., 2020; Wang et al., 2018). *psbs* mutant plants exhibit reduced NPQ capacities and are more susceptible to stress (Acebron et al., 2021). Therefore, the stability of the PsbS protein plays an important role in plant response to abiotic stress. However, little is known about how low temperature regulates PsbS stability to affect energy homeostasis.

The efficient degradation of an entire damaged chloroplast and its specific components is required for efficient photosynthesis and other metabolic reactions under stress conditions (Yang et al., 2019). Recent studies have highlighted that ubiquitination is involved in the degradation of chloroplast-associated proteins, but the exact mechanism remains elusive (Ling et al., 2019). COP9 signalosome (CSN) is an evolutionarily conserved multisubunit protein complex that cleaves the covalently linked Related Ubiquitin 1 (RUB1)/neural precursor cell expressed, developmentally down-regulated 8 (NEDD8) from the CULLIN-RING E3 ubiquitin ligase (CRL) complexes (Lyapina et al., 2001; Schwechheimer et al., 2001), and participates in the ubiquitin-proteasomal pathway of protein degradation (Barth et al., 2016). This catalytic activity is located in the MPN motif of the CSN5 subunit (Cope et al., 2002).

CSN5 is involved in plant development and stress responses by regulating auxin, jasmonic acid, and anthocyanin biosynthesis (Luo et al., 2021; Singh et al., 2019; Shang et al., 2019; Schwechheimer et al., 2001). In Arabidopsis, CSN5 is encoded by two genes, *CSN5A* and *CSN5B* (Tomoda et al., 1999). CSN5A was found to regulate Arabidopsis seed germination by facilitating the protein degradation of the repressor of GA-like 2 (RGL2) and ABA insensitive 5 (ABI5), highlighting the role of CSN5 in protein degradation (Jin et al., 2018). Arabidopsis *csn5a* mutants exhibited an increase in chlorophyll content, CO_2_ assimilation rate, and photosynthetic activity under heat stress (Singh et al., 2019). Given that CSN5A is heavily involved in the physiological processes of plants, it is conceivable that CSN5A has broader roles that include, for example, regulating the stability of chloroplast proteins under stress conditions.

Suppressors of PPI1 locus1 (SP1) and Plant U-Box 4 (PUB4) are the two major E3 ubiquitin ligases reported to be involved in chloroplast ubiquitination, but they seem to only mediate the degradation of the outer membrane of chloroplast components (Ling et al., 2019; Ling and Jarvis, 2016). The question of how intra-chloroplast proteins are ubiquitinated remains uncertain. One possibility is that ubiquitination occurs inside a chloroplast. However, no ubiquitination modifiers were identified in the same. Alternatively, intra-chloroplast proteins were suggested to be modified by E3 ubiquitin ligases, after being transported outside the chloroplast by unknown mechanisms (Li et al., 2022). The nucleus-encoded chloroplast vesiculation (CV) protein is involved in chloroplast degradation that occurs outside an organelle (Wang and Blumwald, 2014). Stress-induced CVs target the chloroplasts and interact with chloroplast proteins such as PsbO. This interaction leads to the formation of CV-containing vesicles (CCVs), mobilizing chloroplast proteins to the central vacuole for degradation (Pan et al., 2023; Wang and Blumwald, 2014). Aside from known substrates, the role of CV in PsbS degradation is unclear. CVs are involved in the efflux of chloroplast proteins, facilitating contact between intra-chloroplast proteins and the cytoplasmic components. However, it is still unclear whether CVs are related to the ubiquitination of chloroplast proteins, or whether CCV can be degraded by the ubiquitin/proteasome system (UPS).

Tomato (*Solanum lycopersicum* L.), cultivated widely as a horticultural crop around the world, is extremely susceptible to low night temperature (LNT) stress in the glasshouse cultivation of vegetables during winter and spring in Northern China, which led to severe reductions in tomato yield and quality (Lu et al., 2021). In this study, we found that LNT stress suppressed the increase of NPQ and led to the degradation of SlPsbS in tomato leaves. SlCSN5A directly interacted with SlPsbS in the cytosol and mediated the ubiquitination and degradation of SlPsbS. SlCV was involved in the translocation of SlPsbS from the chloroplasts, facilitating contact between SlCSN5A and the substrate (SlPsbS). Interestingly, SlCV was also found to physically interact with SlCSN5A to be ubiquitinated and degraded. Further, *SlCSN5*-RNAi and *Slcv* mutant plants exhibited higher tolerance to LNT, while *SlCV* overexpression promoted the degradation of SlPsbS proteins in tomato leaves. On the whole, our study demonstrated that SlCSN5A and SlCV played critical roles in regulating chloroplast protein stability under stress, thereby revealing a potentially new pathway for enhancing plant cold tolerance.

## RESULTS

### Silencing of *SlPsbS* increases tomato plant sensitivity to LNT stress

Under LNT stress, tomato plants gradually wilted (Figure 1A), and the maximum quantum yield of PSII (*F*_v_/*F*_m_) decreased with the prolonged stress time (Figure 1B). Within 8 h of LNT stress, NPQ was significantly induced, while LNT stress for 24 h decreased NPQ (Figure 1C). It has been demonstrated that the most effective excitation quenching requires the presence of the PsbS protein (Welc et al., 2021). Therefore, we further detected the abundance changes of SlPsbS proteins under LNT stress. Similar to NPQ, LNT stress also significantly decreased SlPsbS protein levels in the late stages (Figure 1D).

**Figure 1.**
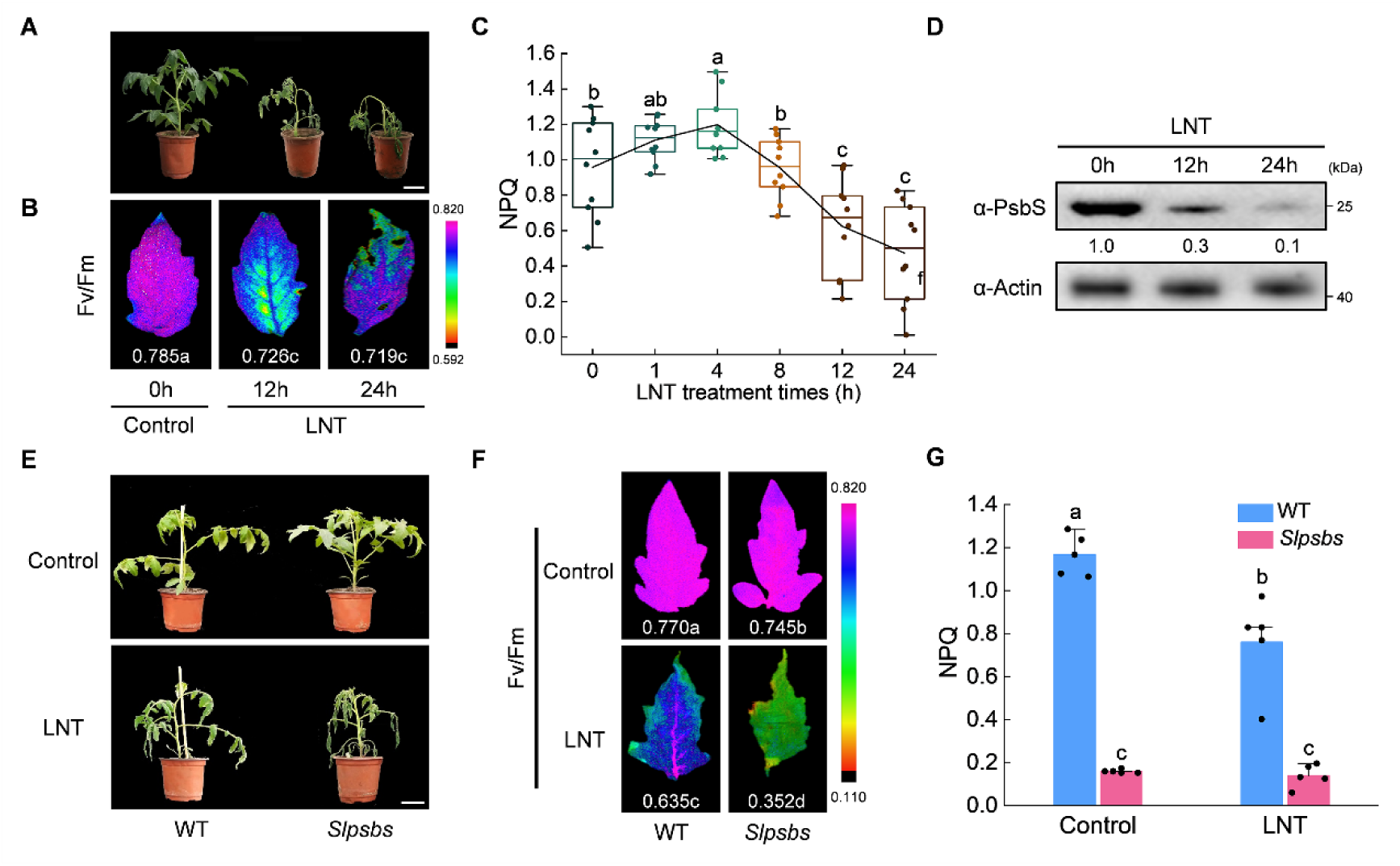
LNT inhibits NPQ and promotes PsbS degradation of tomato leaves. (A) The phenotypes of tomato plants under LNT stress (bar, 5 cm). (B) The maximum quantum yield of PSII (*F*_v_/*F*_m_). (C) Non-photochemical quenching (NPQ). (D) PsbS protein levels were analyzed by immunoblot analysis. Immunoblotting with anti-Actin antibody provided control for equal loading. Intensities of bands were quantified by ImageJ and normalized to Actin and expressed relative to controls. The relative amounts are shown below each lane. The numbers on the right denoted the molecular mass of marker proteins in kiloDaltons. (E) The phenotypes of WT and *Slpsbs* mutant plants before and after LNT treatment (bar, 5 cm). (F) The maximum quantum yield of PSII (*F*_v_/*F*_m_). (G) Non-photochemical quenching (NPQ). Data are the means of five replicates with standard errors shown by vertical bars. Differences among treatments were analyzed by the one-way ANOVA comparison test (P < 0.05). Different letters indicate significant differences among treatments.

To study the role of SlPsbS in tomato plants’ response to LNT stress, we generated *Slpsbs* mutant plants using a CRISPR/Cas9 gene-editing protocol. Western blot showed no accumulation of SlPsbS protein in the *Slpsbs* mutant plants (Figure S1A). Compared to the WT plants, *Slpsbs* mutant plants were found more wilted, with a significant decrease in *F*_v_/*F*_m_ and photosynthetic rate (Pn), and a high accumulation of ROS content under LNT stress (Figures 1E-1F and S1B-S1C). Also, unlike WT plants, which had higher NPQ values under LNT stress, the *Slpsbs* mutant plants showed extremely low NPQ levels under both normal and LNT conditions (Figure 1G). These results indicated that stable SlPsbS contributed to induced NPQ and enhanced tomato LNT tolerance.

### LNT increased ubiquitination levels of SlPsbS

Research revealed that chloroplast proteins can be ubiquitinated and degraded (Li et al., 2022). To study whether the degradation of SlPsbS under LNT stress is related to ubiquitin, we first detected the ubiquitination level of chloroplast proteins under LNT stress. The analysis revealed increased ladder-like smears in chloroplast proteins isolated under LNT conditions (Figure 2A). This suggests that LNT treatment enhances the ubiquitination of chloroplast proteins. We then investigated whether SlPsbS could be ubiquitinated in a transient expression system by fusing SlPsbS to 3x FLAG tag driven by the 35S promoter and transiently converting *N. benthamiana*. After immunoprecipitation with the anti-FLAG beads, ubiquitinated proteins were detected in SlPsbS-3xFLAG samples, but not in 3xFLAG samples (Figure 2B), suggesting that SlPsbS proteins are poly-ubiquitinated in vivo.

**Figure 2.**
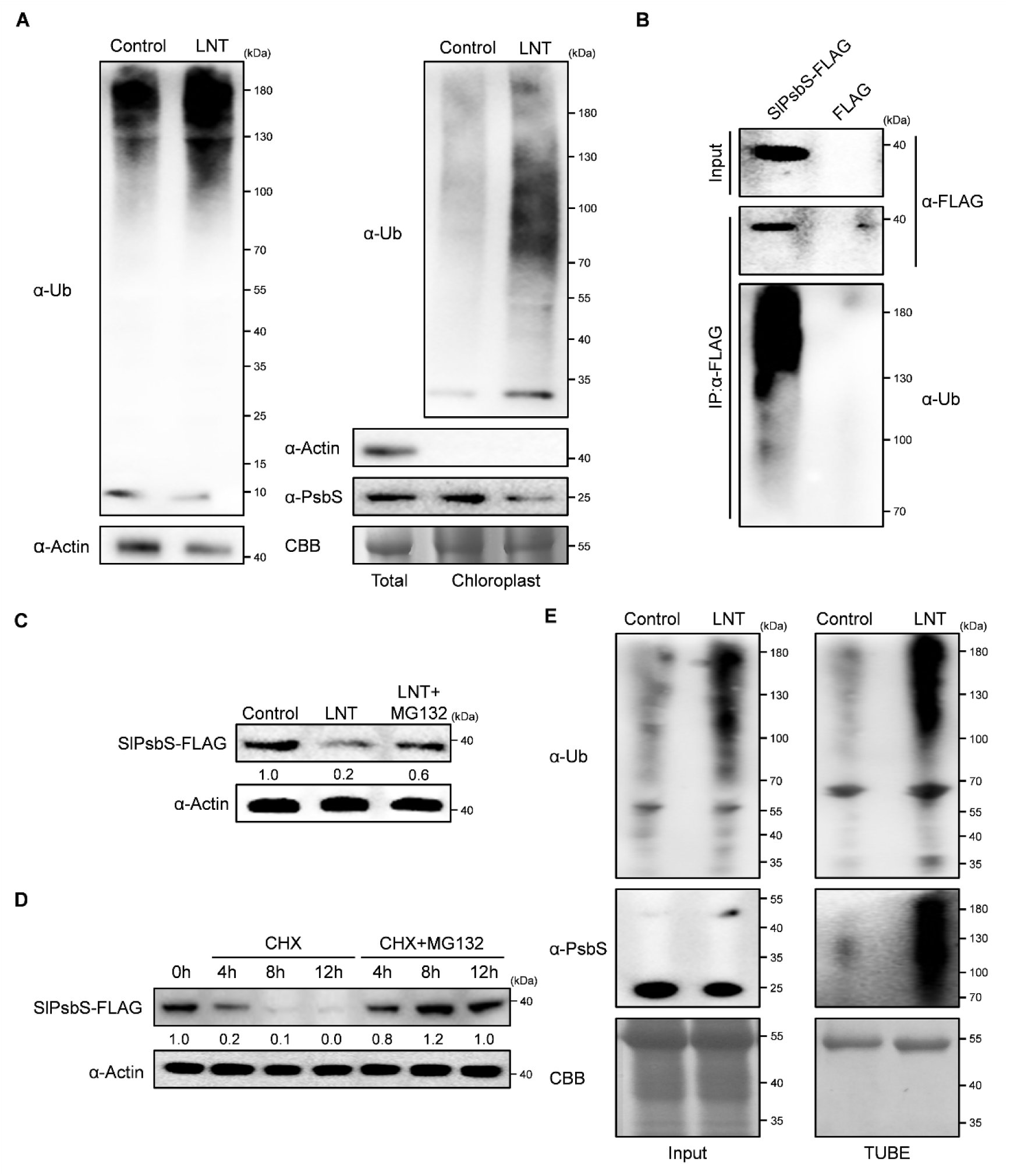
SlPsbS can be ubiquitinated. (A) Ubiquitination levels of total proteins (left panels) and chloroplast proteins (right panels) under control and LNT stress. (B) Ubiquitination assay of SlPsbS. The recombinant plasmids were infected into *N. benthamiana* leaves and immunoprecipitated with anti-FLAG beads. (C) LNT stress leads to degradation of SlPsbS. The SlPsbS-FALG plasmid was transiently into *N. benthamiana* leaves. Four days after infiltration treated with LNT for 24h and 50 μM MG132 (26S proteasome inhibitor) was infiltrated 12 h before sampling. (D) The 26s proteasome system degrades SlPsbS. After SlPsbS-FALG was transfected into *N. benthamiana* leaves for 4 days, they were treated with 50μM CHX (protein synthesis inhibitor in eukaryotes) and CHX+MG132 respectively. The samples were collected at the indicated times and used to determine the level of SlPsbS-FALG protein. (E) Total ubiquitinated proteins were pulled down using tandem ubiquitin-binding entities 2 (TUBE2) from control and LNT-treated tomato leaves. The ubiquitination level of SlPsbS was detected with anti-PsbS antibody. The coomassie brilliant blue staining (CBB) or anti-Actin was a loading control. Intensities of bands were quantified by ImageJ normalized to Actin and expressed relative to controls. The relative amounts are shown below each lane. The numbers on the right denoted the molecular mass of marker proteins in kiloDaltons.

Next, we performed LNT treatment on *N. benthamiana* plants transiently expressing SlPsbS-3xFLAG. As anticipated, the protein abundance of SlPsbS was quickly attenuated under LNT. However, when we treated the plants with MG132, the SlPsbS protein level remained higher under LNT conditions (Figure 2C). Similarly, SlPsbS protein abundance continued to decrease when protein translation was inhibited by CHX, and MG132 treatment promoted the accumulation of SlPsbS in the presence of CHX (Figure 2D). Further, we used a tandem ubiquitin-binding entity (TUBE2) to bind to control and LNT-stressed tomato total protein samples. Immunoblot analysis revealed an increased amount of ubiquitinated SlPsbS pulled down by TUBE2 under LNT compared to controls (Figure 2E). These results suggest that LNT induces the SlPsbS ubiquitination and degradation through 26S proteasome. **SlCSN5A interacts with SlPsbS and mediates its degradation**

It remains uncertain how intra-chloroplast proteins are ubiquitinated. To further analyze the regulatory mechanism of chloroplast protein stability, a yeast two-hybrid (Y2H) screen was performed using SlPsbS as bait. We found that the SlCSN5A could interact with SlPsbS in the Y2H screen. Subsequently, the interaction between SlCSN5A and SlPsbS in yeast cells was verified by the Y2H assay (Figure 3A). Further, we found that the SlPsbS binding site on SlCSN5A was located in MPN rather than in the ICA domain (Figures 3A and S2A). To confirm this *in vivo*, a firefly luciferase (LUC) complementation imaging (LCI) assay was conducted. A strong luminescence signal was detected when constructs containing SlCSN5A fused to the N-terminus of LUC (SlCSN5A-nLUC) and SlPsbS tagged with the C-terminal of LUC (cLUC-SlPsbS) were co-infiltrated into *N. benthamiana* leaves (Figure 3B). In vitro, glutathione S-transferase (GST) pull-down assay found that SlPsbS-HIS was pulled down by SlCSN5A-GST, which further verified that they interact (Figure 3C). Co-immunoprecipitation experiments showed that the protein extracts were subjected to immunoprecipitation using anti-FLAG and that SlCSN5A-GFP were rich in SlPsbS-FLAG immunoprecipitates, which were absent in the negative control (Figure 3D). Next, the effect of SlCSN5A on SlPsbS protein stability was analyzed. After the SlPsbS-FLAG and SlCSN5A-GFP vectors were transiently co-transferred into *N. benthamiana* leaves and treated with CHX, the protein bands of SlPsbS-FLAG were hardly detected by the anti-FLAG antibody. However, the accumulation of SlPsbS-FLAG protein resumed in the presence of MG132 (Figure 3E).

**Figure 3.**
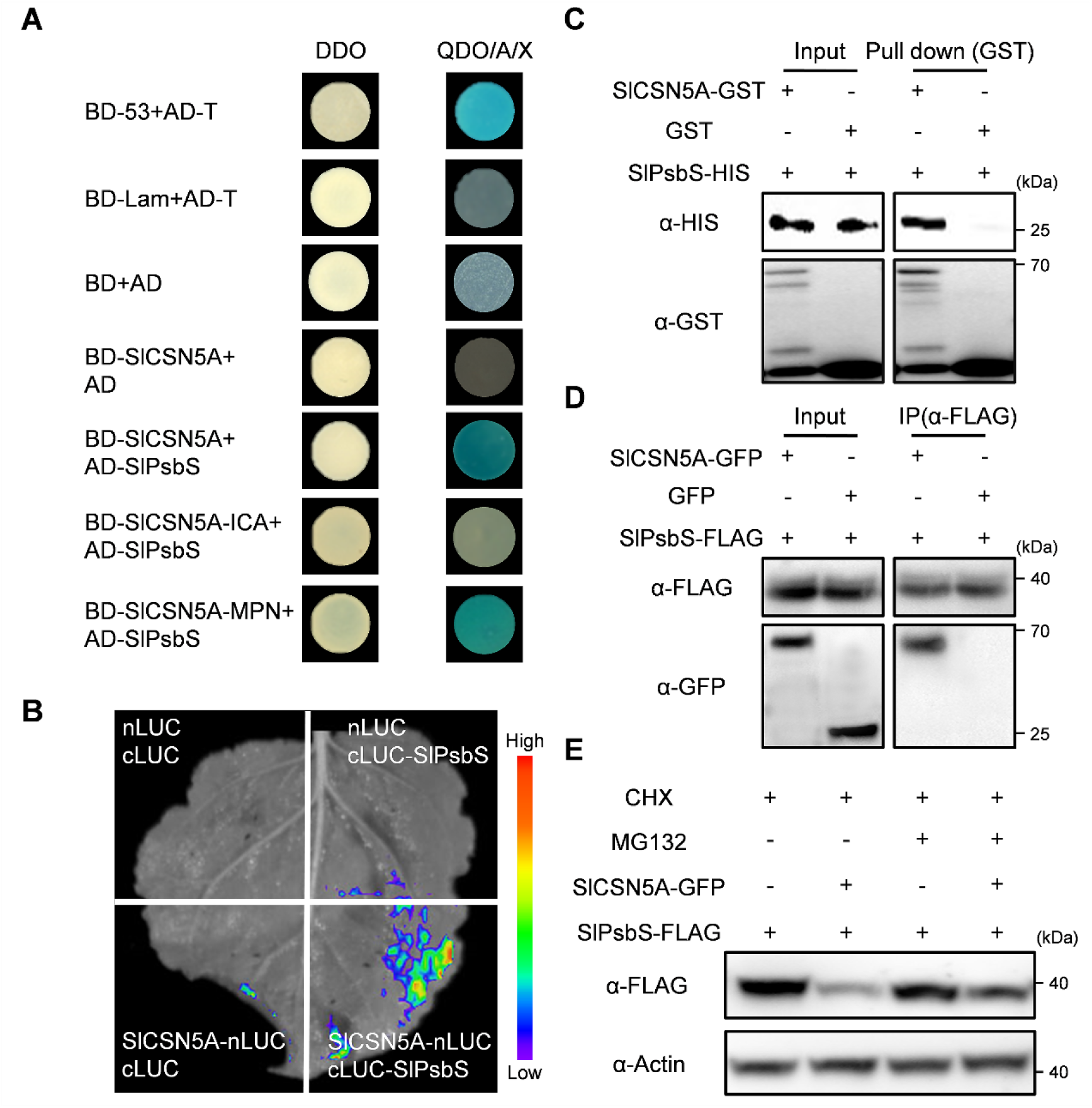
SlCSN5A interacts with SlPsbS and mediates its degradation. (A) Yeast two-hybrid (Y2H) assays showing the interaction between SlCSN5A and SlPsbS. SlCSN5A and SlPsbS were fused to either the DNA binding domain (BD) or the activation domain (AD) of GAL4. DDO, SD medium lacking Trp/Leu; QDO/A/X, SD medium lacking Trp/Leu/His/Ade and containing X-α-gal and aureobasidin A. The empty plasmids were used as controls. Blue color indicates protein interaction. (B) Luciferase complementation imaging assay showing that SlCSN5A interaction with SlPsbS in *N. benthamiana* leaves. *A. tumefaciens* strain EHA105 harboring different constructs was infiltrated into *N. benthamiana* leaves and examined after 72 h. (C) Pull-down assay showing that SlCSN5A interacts with SlPsbS in vitro. The SlCSN5A-GST and SlPsbS-HIS fusion proteins were expressed in *E. coli* and purified. SlCSN5A-GST was bound to GST magnetic beads. The assays were analyzed using immunoblotting with anti-GST and anti-HIS antibodies. (D) Co-Immunoprecipitation assay. SlCSN5A-GFP and SlPsbS-FLAG were cotransformed in *N. benthamiana* leaves. The leaves were collected and extracted total protein, and immunoprecipitation against anti-FLAG beads. Immunoblot analysis with anti-GFP and anti-FLAG antibodies. (E) SlCSN5A promotes SlPsbS degradation by the 26s proteasome. The combination of SlCSN5A-GFP and SlPsbS-FLAG recombinant constructs was co-transformed into *N. benthamiana* leaves. Five days after infiltration, and immunoblotted against anti-FLAG antibody. CHX, protein synthesis inhibitor in eukaryotes. MG132, 26S proteasome inhibitor. The anti-Actin served as a loading control and numbers on the right denoted the molecular mass of marker proteins in kiloDaltons.

### SlCSN5A regulates SlPsbS ubiquitination levels and affects NPQ under LNT stress

LNT stress induced a significant increase in the expression level and protein abundance of SlCSN5A (Figures S2B and S2C). To further study the role of SlCSN5A in the tomato plant response to LNT stress, RNA interference (RNAi) was used to downregulate the expression of *SlCSN5* (including *SlCSN5A* and *SlCSN5B*) in stably transformed plants. It was impossible to independently silence *SlCSN5A* and *SlCSN5B* in tomato plants due to their high sequence similarity (Luo et al., 2021). SlCSN5A gene expression and protein levels were significantly down-regulated in *SlCSN5*-RNAi (*SlCSN5*-RNAi #1, *SlCSN5*-RNAi #3) plants. (Figures S2D and S2E). The *SlCSN5*-RNAi line plants showed decreased sensitivity to LNT stress. Compared to the WT plants, *SlCSN5*-RNAi line plants showed more robust plant phenotypes and higher levels of *F*_v_/*F*_m_, Pn, and NPQ, while ROS content significantly decreased under LNT stress (Figures 4A-4B and S2F-S2H). In addition, *SlCSN5*-RNAi line plants showed higher SlPsbS protein levels compared to the WT plants under LNT stress (Figure 4C). Therefore, SlCSN5A negatively regulates SlPsbS stability and NPQ levels in tomato leaves under LNT.

**Figure 4.**
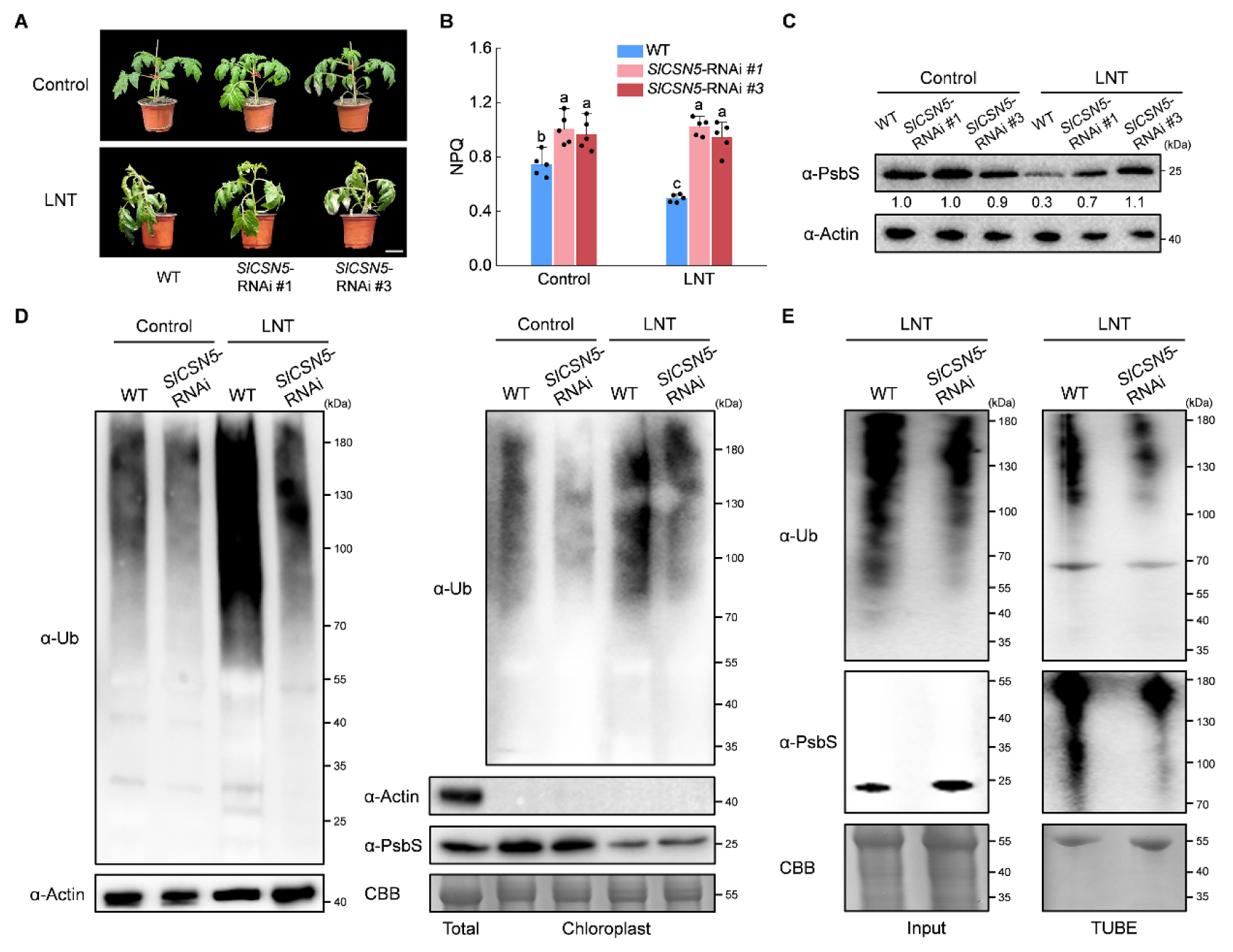
SlCSN5 regulates SlPsbS ubiquitination levels and affects NPQ. (A) The phenotypes of WT and *SlCSN5-*RNAi plants before and after LNT treatment (bar, 5 cm). (B) Non-photochemical quenching (NPQ). Data are the means of five replicates with standard errors shown by vertical bars. Differences among treatments were analyzed by the one-way ANOVA comparison test (P < 0.05). Different letters indicate significant differences among treatments. (C) PsbS protein levels were analyzed by immunoblot analysis in WT and *SlCSN5-*RNAi plants before and after LNT treatment. Intensities of bands were quantified by ImageJ normalized to Actin and expressed relative to controls. The relative amounts are shown below each lane. (D) Ubiquitination levels of total proteins (left panels) and chloroplast proteins (right panels) in WT and *SlCSN5-*RNAi plants under control and LNT stress. (E) Total ubiquitinated proteins were pulled down using tandem ubiquitin-binding entities 2 (TUBE2) from WT and *SlCSN5-*RNAi plants under control and LNT stress. The ubiquitination level of SlPsbS was detected with anti-PsbS antibody. Immunoblotting with coomassie brilliant blue staining (CBB) or anti-Actin antibody provided control for equal loading. The numbers on the right denoted the molecular mass of marker proteins in kiloDaltons.

Next, we examined the ubiquitination levels of total and chloroplast proteins in *SlCSN5*-RNAi line and WT plants. LNT promoted the accumulation of ubiquitinated proteins in WT compared to controls, whereas fewer high-molecular weight smears were detected in *SlCSN5*-RNAi line regardless of the temperature conditions. Similarly, *SlCSN5*-RNAi line also accumulated less ubiquitinated proteins than WT in their chloroplasts (Figure 4D). We further used TUBE2 to capture all ubiquitinated proteins in the LNT-treated *SlCSN5*-RNAi line and WT plants and measured the levels of captured *SlPsbS* proteins. Immunoblot analysis showed that the amount of ubiquitin-conjugated forms of SlPsbS in the *SlCSN5*-RNAi line was significantly reduced compared with WT (Figure 4E). To detect the ubiquitin ligase activity of SlCSN5A, we performed in vitro ubiquitination assays. No polyubiquitination conjugation was observed in the SlCSN5A-GST purified proteins, suggesting that SlCSN5A is not a functional E3 ligase (Figure S3). Together, these results support a role for SlCSN5A in the ubiquitination and degradation of SlPsbS in the LNT.

### SlCV promotes interaction between SlPsbS and SlCSN5A

Subcellular localization analysis showed that SlPsbS was located in the chloroplast (Figure S4A). However, green fluorescence signals of transiently expressed SlCSN5A-GFP in wild tobacco leaves coincided with the cytosol localized HPR2-mCherry signal (Figure S4B), indicating the CSN5A is present in the cytosol (Mo et al., 2021). As chloroplasts are organelles surrounded by double membranes, it is unclear if or how SlPsbS is retranslocated from the chloroplasts to come into contact with SlCSN5A and UPS complexes in the cytosol. Wang and Blumwald (2014) showed that the CV protein induced the formation of CCVs, which were released from chloroplasts mobilized to the vacuole through cytosol for proteolysis. Co-translocation of CV with chloroplast proteins appears to allow for contacts of intra-chloroplast proteins with cytosol components. Therefore, we speculated that CCVs containing SlPsbS were excreted from the chloroplasts, and in the cytoplasm, CCVs contributed to the contact of SlPsbS with SlCSN5A. To prove this, the expression level of *SlCV* was examined under LNT stress. The *SlCV* expression level increased significantly with the aggravation of stress, which was up-regulated 356-fold under 24-h stress conditions (Figure S5A). Similarly, SlCV protein abundance also significantly accumulated under LNT stress (Figure S5B). We further analyzed the subcellular localization of SlCV. Using a confocal laser scanning microscope, it was observed that the green fluorescence signal of SlCV-GFP driven by the CaMV35S promoter coincided with the autofluorescence of chloroplasts, indicating that SlCV was located within the chloroplasts (Figure S5C). Interestingly, after LNT treatment of *N. benthamiana* plants expressing SlCV-GFP, green fluorescence signals were found outside the chloroplasts in some unknown compartments (Figure S5D).

To determine the role of SlCV in SlPsbS degradation, the interaction between SlCV and SlPsbS was analyzed by Y2H. The results showed that SlCV interacted with SlPsbS in yeast cells (Figure 5A). We further confirmed the interaction between SlCV and SlPsbS in LCI assay (Figure 5B) and pull-down assay (Figure 5C). In co-immunoprecipitation analysis, SlCV-GFP was co-immunoprecipitated with SlPsbS-FLAG through immunoblotting using the anti-FLAG beads (Figure 5D). In addition, SlCV-GFP and SlPsbS-mCherry were transiently co-expressed in N. benthamiana leaves. SlCV-GFP and SlPsbS-mCherry were co-localized in chloroplasts under control treatment while overlapping fluorescence signals were detected outside the chloroplasts under LNT treatment (Figure 5E). This raises the question of whether this phenomenon is related to the interaction between SlPsbS and SlCSN5A. To address this, a BiFC assay was conducted to determine the specific location of SlCSN5A and SlPsbS interaction in planta. Co-infiltration of SlCSN5A-nYFP and cYFP-SlPsbS in *N. benthamiana* leaves resulted in the generation of a YFP fluorescence signal in the cytosol, which was further enhanced under LNT treatment. Interestingly, the addition of SlCV-3FLAG to the BiFC system significantly increased the YFP fluorescence signal, particularly under LNT conditions (Figures 5F and 5G). These findings suggest that SlCV plays a role in facilitating the interaction between SlPsbS and SlCSN5A in the cytosol.

**Figure 5.**
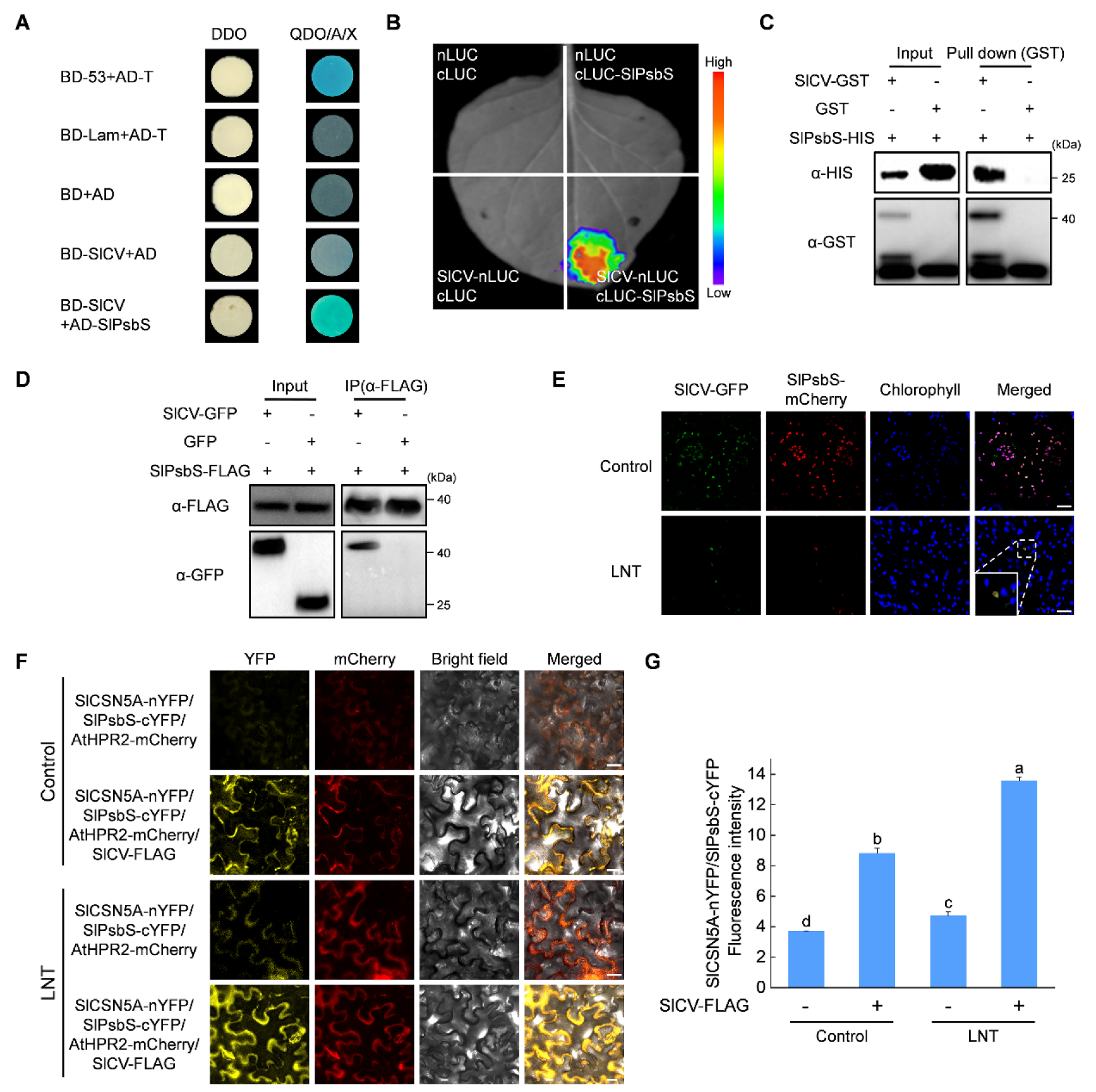
SlCV interacts with SlPsbS. (A) Yeast two-hybrid (Y2H) assays showing the interaction between SlCV and SlPsbS. SlCV and SlPsbS were fused to either the DNA binding domain (BD) or the activation domain (AD) of GAL4. DDO, SD medium lacking Trp/Leu; QDO/A/X, SD medium lacking Trp/Leu/His/Ade and containing X-α-gal and aureobasidin A. The empty plasmids were used as controls. Blue color indicates protein interaction. (B) Luciferase complementation imaging assay showing that SlCV interaction with SlPsbS in *N. benthamiana* leaves. *A. tumefaciens* strain EHA105 harboring different constructs was infiltrated into *N. benthamiana* leaves and examined after 72 h. (C) Pull-down assay showing that SlCV interacts with SlPsbS in vitro. The SlCV-GST and SlPsbS-HIS fusion proteins were expressed in *E. coli* and purified. SlCV-GST was bound to GST magnetic beads. The assays were analyzed using immunoblotting with anti-GST and anti-HIS antibodies. SlCV and SlPsbS interaction in a coimmunoprecipitation (Co-IP) assay. SlCV-GFP and SlPsbS-FLAG were co-transformed in *N. benthamiana* leaves. The proteins were immunoprecipitation against anti-FLAG beads. Immunoblot analysis with anti-GFP and anti-FLAG antibodies. The numbers on the right denoted the molecular mass of marker proteins in kiloDaltons. (E) Colocalization analysis of SlCV-GFP and SlPsbS-mCherry. Two days after SlCV-GFP and SlPsbS-mCherry co-transformed *N. benthamiana* leaves, they were treated with LNT for 24h. The fluorescence signal was monitored with a confocal laser scanning microscope (bar, 25 μm). (F) SlCV promotes interaction between SlPsbS and SlCSN5A. SlCSN5A-nYFP and SlPsbS-cYFP were co-infiltrated into *N. benthamiana* leaves and expressed for 72 h and BIFC fluorescence signals were detected in the presence or absence of SlCV-FLAG under control or LNT conditions. HPR2-mCherry was used as a cytosol marker (bar, 25 μm). (G) SlCSN5A-nYFP and SlPsbS-cYFP fluorescence intensity. All data were captured at the same exposure times, contrast settings, and intensity for measurement of fluorescence intensity. Data are the means of four replicates with standard errors shown by vertical bars. Differences among treatments were analyzed by the one-way ANOVA comparison test (P < 0.05). Different letters indicate significant differences among treatments.

### SlCV inhibits NPQ production and promotes PsbS degradation in tomato leaves under LNT stress

*SlCV* overexpression (*SlCV*-OE #91, *SlCV*-OE #92) transgenic tomato plants were obtained under the control of the CaMV35S constitutive promoter. Compared to WT, the *SlCV* expression levels and SlCV protein levels showed a significant increase in *SlCV*-OE #91 and *SlCV*-OE #92 plants (Figures S6A and S6B). Under normal conditions, *SlCV*-OE #91 and *SlCV*-OE #92 plants were significantly shorter compared to the WT plants (Figure 6A). In addition, *SlCV*-OE #91 and *SlCV*-OE #92 showed increased sensitivity to LNT stress, as evidenced by severe plant wilting, significant decrease in *F*_v_/*F*_m_, Pn, and massive accumulation of ROS levels (Figures S6C-S6E). Unlike higher NPQ values in the WT plants, *SlCV*-OE #91 and *SlCV*-OE #92 plants showed lower NPQ levels (Figure 6B). Further, *SlCV*-OE #91 and *SlCV*-OE #92 plants showed lower levels of SlPsbS protein abundance than WT, under control and LNT conditions (Figure 6C).

**Figure 6.**
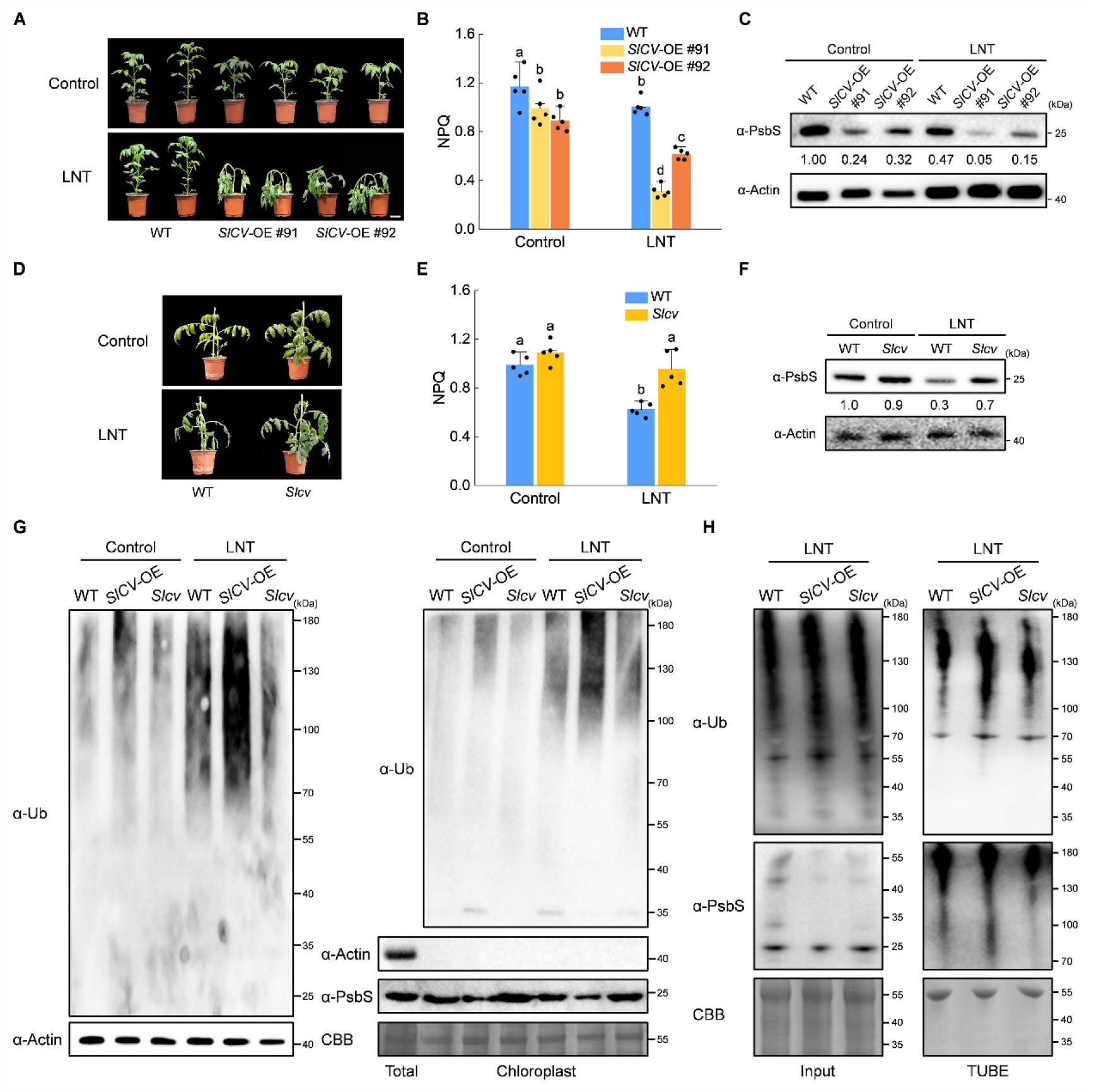
SlCV inhibits NPQ production in tomato leaves under LNT stress. (A) The phenotypes of WT and *SlCV* overexpressing plants before and after LNT treatment (bar, 5 cm). (B) Non-photochemical quenching (NPQ). (C) PsbS protein levels of WT and *SlCV* overexpression plants before and after LNT treatment. The numbers on the right denoted the molecular mass of marker proteins in kiloDaltons. (D) The phenotypes of WT and *Slcv* mutant plants before and after LNT treatment (bar, 5 cm). (D) Non-photochemical quenching (NPQ). Data are the means of five replicates with standard errors shown by vertical bars. Differences among treatments were analyzed by the one-way ANOVA comparison test (P < 0.05). Different letters indicate significant differences among treatments. (F) PsbS protein levels were analyzed by immunoblot analysis in WT and *Slcv* plants before and after LNT treatment. Immunoblotting with anti-Actin antibody provided control for equal loading. Intensities of bands were quantified by ImageJ normalized to Actin and expressed relative to controls. The relative amounts are shown below each lane. The numbers on the right denoted the molecular mass of marker proteins in kiloDaltons. (G) Ubiquitination levels of total proteins (left panels) and chloroplast proteins (right panels) in WT, *SlCV* overexpression and *Slcv* mutant plants under control and LNT stress. (H) Total ubiquitinated proteins were pulled down using tandem ubiquitin-binding entities 2 (TUBE2) from WT, *SlCV* overexpression and *Slcv* mutant plants under control and LNT stress. The ubiquitination level of SlPsbS was detected with anti-PsbS antibody. Immunoblotting with coomassie brilliant blue staining (CBB) or anti-Actin antibody provided control for equal loading. The numbers on the right denoted the molecular mass of marker proteins in kiloDaltons.

*Slcv* mutant plants were obtained using the CRISPR/Cas9 gene-editing system to further study the role of SlCV in LNT stress (Figure S7A). Under LNT stress, *Slcv* mutant plants maintained normal plant morphology and lower ROS levels compared to the WT plants (Figures 6D and S7B). *Slcv* mutants also exhibited an increase in *F*_v_/*F*_m_ and Pn under LNT conditions (Figures S7C and S7D). Meanwhile, LNT induced higher NPQ values and SlPsbS protein abundance in *Slcv* mutants compared to the WT plants (Figures 6E and 6F).

### SlCV affects ubiquitination levels of chloroplast proteins

Next, we analyzed whether SlCV would affect the ubiquitination levels of chloroplast proteins and SlPsbS. First, we detected ubiquitination levels of total and chloroplast protein in SlCV-OE and *Slcv* mutant plants under control and LNT conditions with anti-Ub antibodies. We found that LNT treatment promoted the accumulation of ubiquitinated proteins in WT, whereas more ubiquitinated products were detected in total protein and chloroplast fractions of SlCV-OE plants than in WT. In contrast, only weak high molecular weight smears are present in *Slcv* mutants (Figure 6G). Further, we used TUBE2 to enrich all ubiquitinated proteins in SlCV-OE and *Slcv* mutant plants under LNT to evaluate the ubiquitination level of SlPsbS. More ubiquitylated forms of SlPsbS were detected in SlCV-OE plants compared to WT plants, but only small amounts of ubiquitylated SlPsbS pulled down by TUBE2 were observed in *Slcv* plants (Figure 6H). The results mentioned above suggest that the degradation of SlPsbS by SlCV may also involve ubiquitination.

### SlCSN5A interacts with SlCV

To understand the relationship among SlCV, SlCSN5A, and SlPsbS, a possible protein-protein interaction between SlCV and SlCSN5A was investigated. We demonstrated that SlCV interacted with SlCSN5A in both yeast cells and plants using Y2H and LCI assays (Figure S8A and S8B). Further, the SlCV binding site on SlCSN5A was found in MPN rather than in the ICA domain (Figure S8A). We then purified SlCV-GST and SlCSN5A-HIS fusion proteins and performed a pull-down assay to confirm the interaction between SlCV and SlCSN5A (Figure S8C). In vivo, SlCSN5A-GFP and SlCV-FLAG were co-transformed into *N. benthamiana* leaves and immunoprecipitated using the anti-FLAG antibody. SlCSN5A-GFP protein was co-immunoprecipitated with SlPsbS-FLAG but was absent in the negative control (Figure S8D). In the BIFC analysis, SlCV fused with nYFP interacted with SlCSN5A fused with cYFP, generating YFP fluorescence in the cytosol (Figure S8E).

The interaction between SlCSN5A and SlCV prompted us to hypothesize that CV may also be a substrate for ubiquitination. As anticipated, after immunoprecipitation with the anti-FLAG beads, we observed the presence of high molecular weight smears in *N. benthamiana* leaves expressing SlCV-FLAG, but not in FLAG alone (Figure S9A), suggesting that SlCV proteins are poly-ubiquitinated in vivo. We conducted further analysis on the impact of SlCSN5A on SlCV stability. In the presence of CHX, there was minimal SlCV accumulation observed when *N. benthamiana* leaves were co-infected with SlCSN5A-GFP and SlCV-FLAG. However, upon treatment with MG132, the SlCV stability improved (Figure S9B). These findings indicate that SlCSN5A plays a role in regulating SlCV stability through the ubiquitin-proteasome system.

### SlCSN5A is required for SlCV-mediated SlPsbS degradation and sensitivity to LNT stress

To examine the function of SlCSN5A and SlCV in the ubiquitination and degradation of SlPsbS, the *SlCSN5A* gene was silenced in *SlCV*-OE #91 and *SlCV*-OE #92 plants (Figures S10A and S10B). Under LNT stress, *F*_v_/*F*_m_, Pn, NPQ, and SlPsbS protein abundance of tobacco rattle virus (TRV)-based vectors (pTRV) in *SlCV*-OE #91 and *SlCV*-OE #92 plants were significantly lower compared to pTRV in the WT plants. However, silencing of *SlCSN5A* in *SlCV*-OE plants significantly improved the tolerance of *SlCV*-OE plants to LNT stress (Figure 7). Compared to the pTRV plants with *SlCV*-OE #91 and *SlCV*-OE #92 backgrounds, pTRV-*SlCSN5A* plants with *SlCV*-OE #91 and *SlCV*-OE #92 backgrounds showed higher levels of *F*_v_/*F*_m_, Pn, and NPQ under LNT stress (Figures 7B-7D). Analysis of SlPsbS protein levels found that silencing of the *SlCSN5A* gene could delay the degradation of SlPsbS protein in *SlCV*-OE plants under LNT stress (Figure 7E).

**Figure 7.**
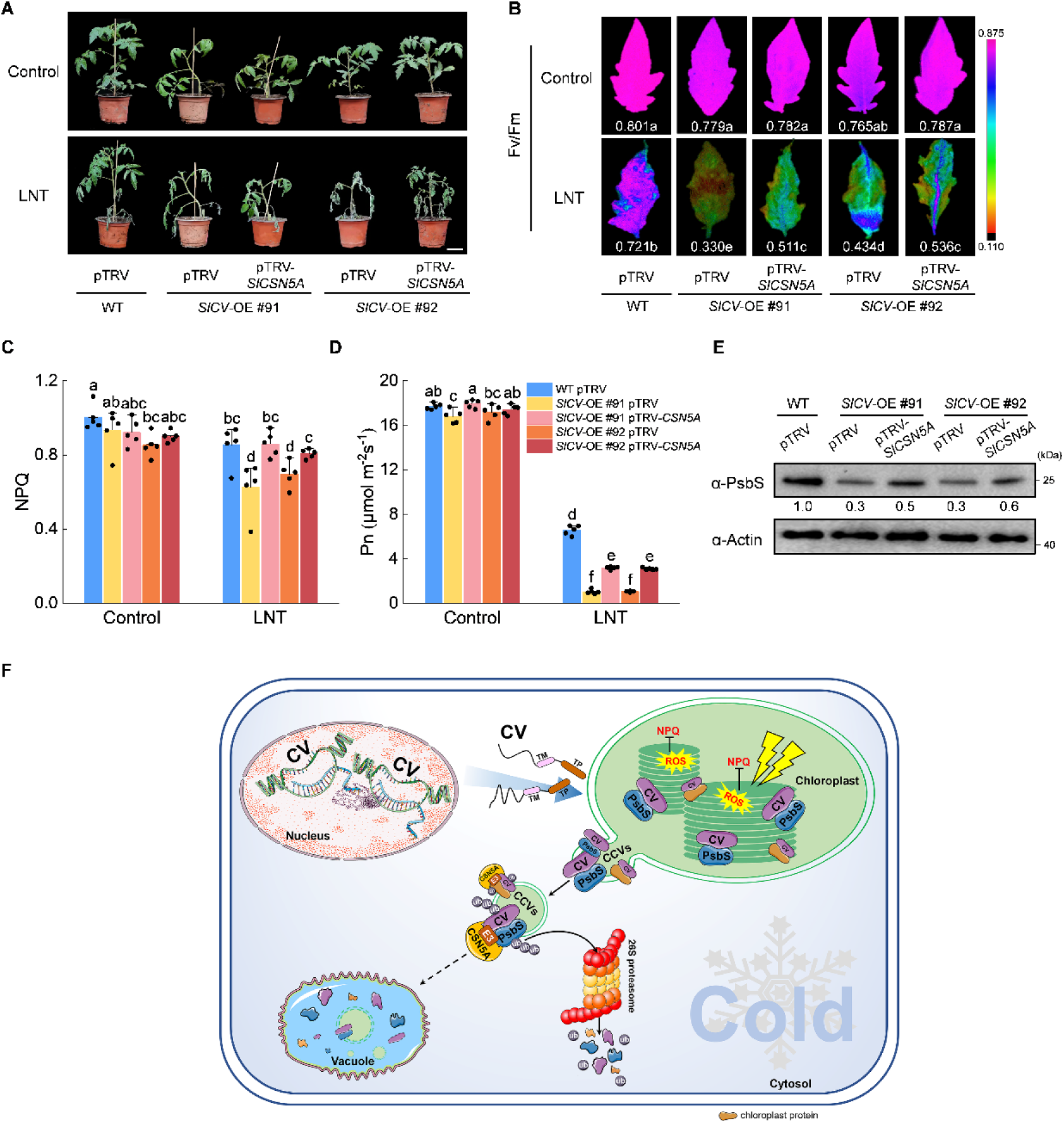
SlCSN5 regulates the stability of SlPsbS in *SlCV* overexpressing plants. (A) The phenotypes in tomato plants when silenced (pTRV-*SlCSN5A*) in *SlCV* -OE plants and nonsilenced *SlCSN5A* (pTRV) in WT and *SlCV* -OE plants before and after LNT treatment (bar, 5 cm). (B) The maximum quantum yield of PSII (*F*_v_/*F*_m_). (C) Non-photochemical quenching (NPQ). (D) Net photosynthetic rate (Pn). (E) PsbS protein levels of silenced (pTRV-*SlCSN5A*) in *SlCV*-OE plants and nonsilenced *SlCSN5A* (pTRV) in WT and *SlCV* -OE plants after LNT treatment. Immunoblotting with anti-Actin antibody provided control for equal loading. Intensities of bands were quantified by ImageJ and normalized to Actin and expressed relative to controls. The relative amounts are shown below each lane. The numbers on the right denoted the molecular mass of marker proteins in kiloDaltons. Data are the means of five replicates with standard errors shown by vertical bars. Differences among treatments were analyzed by the one-way ANOVA comparison test (P < 0.05). Different letters indicate significant differences among treatments. (F) Schematic presentation of SlCSN5 regulates SlPsbS protein stability through the ubiquitin-proteasome system under LNT stress. Cold stress activates *SlCV*, SlCV protein targets chloroplast to interact with SlPsbS and induces the formation of CCVs containing chloroplast proteins (SlPsbS), and then CCVs was released from the chloroplast. In the cytosol, SlCV and SlPsbS contact SlCSN5A and the unknown E3 ubiquitin ligase through the extramembrane region, which is then ubiquitinated and degraded by the UPS. The SlPsbS protein abundance decreased will lead to low levels of NPQ value and ROS large accumulation, which exacerbates photosystem photodamage and reduces plant stress tolerance.

## DISCUSSION

Chloroplasts are highly sensitive to stress conditions. The degradation of photosynthetic membrane-coupled proteins and the inhibition of protein synthesis at low temperatures damage the integrity of photosynthetic membranes, thereby impairing light reactions (Gan et al., 2019). Similarly, we show here that LNT stress leads to a significant decrease in *F*_v_/*F*_m_ (Figure 1B). This reduces photosynthetic efficiency, resulting in severe photoinhibition of the photosystem (Lu et al., 2020a; Lu et al., 2021). However, plants have developed a mechanism to dissipate excess light energy in the form of heat through light-harvesting complexes of the photosystem to prevent ROS damage. This is one of the most important, fast photoprotective mechanisms, with the process being called NPQ (Ruban, 2018). The protein PsbS was identified as a key factor responsible for NPQ activation in plants (Krishnan-Schmieden et al., 2021). In this study, NPQ was rapidly induced by LNT and decreased in the late stage of stress, which might be related to the reduction of SlPsbS protein abundance (Figures 1C and 1D). *SlpsbS* mutants showed a slower increase in NPQ values and were highly susceptible to LNT stress (Figure 1), which was consistent with the findings on *Chlamydomonas reinhardtii* and Arabidopsis (Acebron et al., 2021; Redekop et al., 2020). These results suggested that NPQ mediated by PsbS played an important role in photoprotection under stress. Consequently, increasing the stability of PsbS protein may be an effective way to improve plant stress tolerance.

The UPS is a major regulatory system targeting misfolded or unnecessary protein degradation, which plays a crucial role in their growth and development (Xu and Xue, 2019). Ubiquitin-dependent modification of proteins located on the chloroplast’s outer membrane and photodamaged chloroplasts was a pivotal event (Jeran et al., 2021; Ling et al., 2012). Ubiquitome studies showed that chloroplast proteins accounted for a large portion of the total pool of ubiquitinated proteins in various plant species (Rosnoblet et al., 2021; Lu et al., 2019). Recent reports revealed ubiquitin modification’s biological and biochemical functions in controlling the degradation of chloroplast-resident proteins (Li et al., 2022; Sun et al., 2022). Consistent with this notion, we found that LNT stress promoted the enrichment of ubiquitinated proteins in chloroplasts and SlPsbS proteins are poly-ubiquitinated in vivo (Figure 2). Further, SlCSN5A was observed as a novel regulator mediating the ubiquitination and degradation of intra-chloroplast proteins, providing new insights into the regulatory mechanisms underlying chloroplast protein quality control.

CSN is an evolutionarily conserved protein complex (Singh et al., 2019) and best known as a regulator of the superfamily of CRLs, which catalyze a key step in protein ubiquitination and degradation through UPS (Lyapina et al., 2001; Schwechheimer et al., 2001). In tomatoes, SlCSN5 was found to regulate ascorbic acid and anthocyanin biosynthesis by affecting the stability of SlZF3 and B-box protein 20 (SlBBX20) (Luo et al., 2021; Li et al., 2018). Following heat stress, Arabidopsis *csn5a* mutants exhibited an increase in CO_2_ assimilation rate and photosynthetic activity, suggesting that CSN5A is related to photosynthesis or chloroplasts (Singh et al., 2019). In the present study, *SlCSN5* deficiency enhanced the SlPsbS protein abundance and NPQ value in tomato leaves under LNT stress and reduced the sensitivity of plants to stress (Figure 4). CSN5A is highly conserved between plant and animal kingdoms. It regulates UPS to specifically guide ubiquitylated proteins to the 26S proteasome for degradation in plants, animals, and fungi (Schwechheimer, 2001). The new results reported here provide evidence that CSN5A was involved in the chloroplast protein turnover, revealing a different pathway for chloroplast protein degradation compared to previous studies and extending the existing models of CSN5A function.

UPS-mediated protein degradation requires physical interaction between the degradation machinery and its substrates. In our study, SlCSN5A directly interacted with SlPsbS under both *in vivo* and *in vitro* conditions (Figure 3). However, available studies indicated the absence of E3 ubiquitin ligase activity in CSN5A (Figure S3). Hence, the E3 ubiquitin ligase-mediated modification of chloroplast proteins and its association with CSN5A is still largely unknown. Importantly, PsbS is constitutively present in the thylakoid membranes, with no evidence of its association with the chloroplast envelope (Daskalakis and Papadatos, 2017; Li et al., 2000). Numerous studies have indicated the localization of CSN5 in both the nucleus and cytosol (Mo et al., 2021; Luo et al., 2021; Gusmaroli et al., 2004). Our results showed that SlPsbS interacted with SlCSN5A in the cytosol (Figure 5F). However, the translocation of substrates from the chloroplasts and their contact with CSN5A in the cytosol are yet to be resolved.

Studies have shown that abiotic stress activates *CV* transcription (Umnajkitikorn et al., 2020; Sade et al., 2018; Wang and Blumwald, 2014). Similarly, our results revealed that LNT stress induced high expression of *SlCV* and promoted SlCV translocation from the chloroplasts (Figure S5). In Arabidopsis, CV was found to destabilize the chloroplasts and induce the formation of CCVs. CCVs carry stromal proteins, envelope membrane proteins, and thylakoid membrane proteins, which are eventually released from the chloroplasts to the cytosol (Wang and Blumwald, 2014). Hence, we speculated that SlCV represented a bridge linking SlPsbS and SlCSN5A and examined the relationship among SlCV, SlPsbS, and SlCSN5A. It was found that during LNT stress, SlCV directly targets and destroys SlPsbS. Subsequently, CCVs containing damaged SlPsbS are released from chloroplasts, leading to the contact of SlPsbS with cytosolic components (Figures 5 and 6). Additionally, SlSlCV enhanced the interaction strength between SlPsbS and SlCSN5A, and the BIFC signal of SlCV fused with SlCSN5A was also observed in the cytosol (Figures 5F-5G and S8E). These data strongly suggested that SlCV could translocate along with SlPsbS from the chloroplasts to the cytosol, contributing to the contact between the substrate (chloroplast protein), SlCSN5A, and the unknown E3 ubiquitin ligase. Besides SlPsbS, SlCV can also interact with multiple chloroplast proteins. Interestingly, SlCSN5A appeared to regulate the ubiquitination level of every SlCV-interacting intra-chloroplast protein (data not shown). These results further confirmed that SlCV and SlCSN5A played critical roles in the ubiquitination and degradation of intra-chloroplast proteins and that the SlCV and SlCSN5A pathways appeared to have a broad impact on intra-chloroplast proteins stability.

CCVs can carry CV and chloroplast proteins out of chloroplasts and aggregate in the cytosols of the mesophyll cells in true leaves (Wang and Blumwald, 2014). In the present study, SlCSN5A regulated the ubiquitination levels of SlPsbS in the cytosol and promoted SlCV degradation through the ubiquitin-proteasome system (Figures 4 and S8). This suggested that SlCSN5A is involved in the ubiquitination and degradation of CCVs containing SlCV and chloroplast proteins, which is a new degradation mechanism of CCVs. CV proteins are primarily associated with thylakoid membranes and envelope membranes before the formation of CCVs, while CCVs are considered to be a vesicle structure formed directly from the chloroplast membranes disrupted by CV (Pan et al., 2023; Wang and Blumwald, 2014). The presence of CV homologs in all plant species contains a conserved transmembrane domain (Wang and Blumwald, 2014). The association of SlCV and SlPsbS with SlCSN5A in CCVs can be related to its predicted transmembrane domain. As observed in Arabidopsis, CCV formed by CV and chloroplast proteins can translocate into the vacuole (Wang and Blumwald, 2014). Therefore SlCV and SlPsbS may also be mobilized to the vacuole for proteolysis. Recent research showed that, aside from the 26S proteasome pathway, two main components of ESCRT-I, namely vacuolar protein sorting 23A (VPS23A) and FYVE1/FYVE domain protein are required for endosomal sorting 1 (FREE1) to recognize PYL4 ubiquitinated by the E3 ligase RSL1 and promote PYL4 trafficking into the vacuoles for degradation (Belda-Palazon et al., 2016; Yu et al., 2016; Bueso et al., 2014). Thus, the ubiquitination and degradation of SlCV and SlPsbS mediated by SlCSN5A were not contradictory to the vacuolar proteolysis of CCVs. On the other hand, the vacuolar proteolysis of CCVs could be related to ubiquitination.

CV plays a key role in the destabilization of the photosynthetic apparatus during abiotic stress. *CV* overexpression disrupts ROS homeostasis, resulting in excessive accumulation of ROS and decreased stability in both chloroplasts and chloroplast proteins (Yu et al., 2022; Wang and Blumwald, 2014). Silencing of *CV* delays abiotic stress-induced chloroplast turnover and enhances source fitness (i.e., carbon and nitrogen assimilation) and photorespiration (Umnajkitikorn et al., 2020; Sade et al., 2018; Wang and Blumwald, 2014). Under LNT stress, *SlCV* overexpression led to a significant decrease in NPQ and accelerated degradation of SlPsbS, whereas *Slcv* mutants showed higher NPQ values (Figure 6). This indicated the role of CV in regulating NPQ levels and in photoprotection responses to stress. Under heat stress, *CSN5*-silenced Arabidopsis plants increased heat tolerance by modulating auxin signaling (Singh et al., 2019). Similarly, *SlCSN5*-RNAi plants also exhibited enhanced resistance and SlPsbS protein stability under LNT stress (Figure 4). This suggested that the degradation of chloroplast proteins by SlCSN5A was more destructive compared to the ubiquitination and degradation of SlCV and that SlCV might play more the role of a linker between SlCSN5A and chloroplast protein. It is worth noting that the silencing of *SlCSN5A* in tomato CV-OE plants partially rescued the LNT-sensitive phenotype of CV-OE plants and promoted SlPsbS protein accumulation (Figure 7). These genetic observations indicated that the SlCV-induced stress sensitivity and chloroplast protein instability of tomato plants were partially dependent on SlCSN5A.

Based on the current and earlier findings, we constructed a model to describe the impact of SlCV and SlCSN5A on tomato LNT tolerance through the regulation of SlPsbS protein stability. LNT activated *SlCV* expression and the SlCV protein targeted the chloroplast to interact with the SlPsbS and induce the formation of CCVs containing chloroplast proteins. CCVs were released from the chloroplast. In the cytosol, CCVs contacted SlCSN5A and the unknown E3 ubiquitin ligase, which was then ubiquitinated and degraded by UPS. A decrease in SlPsbS protein abundance led to low NPQ values, which exacerbated photosystem photodamage and reduced plant stress tolerance (Figure 7F). On the whole, our work established a potentially new avenue to regulate chloroplast protein stability under stress conditions.

## MATERIALS AND METHODS

### Plant material and growth conditions

*Slpsbs* and *Slcv* mutants were obtained by CRISPR/Cas9 technology according to the method of Shimatani et al. (2017). To generate transgenic *SlCV*-overexpression (*SlCV*-OE) tomato plants, the full-length coding region of *SlCV* (without the stop codon) was PCR-amplified and inserted into the pHSN6A01 vector (containing a 4×minimal VP16 activation domain). *SlCSN5*-RNAi plants were generously provided by Taotao Wang (Huazhong Agriculture University, China) (Luo et al., 2021). TRV-based vectors (pTRV1/2) were used for virus-induced gene silencing (VIGS) of the *SlCSN5A* gene in tomato plants following the method of Singh et al. (2022). WT tomato plants used in this study were of the ‘Ailsa Craig’ ecotype, while the transgenic tomato plants were obtained by *A. tumefaciens*-mediated transformation. The *Slpsbs* and *Slcv* mutants were identified by PCR and confirmed by DNA sequencing. *SlCV*-OE, *SlCSN5*-RNAi, and pTRV-*SlCSN5A* plants were quantified using qRT-PCR. Tomato seeds were germinated and grown in pots under cool-white fluorescent light (600 μmol m^−2^ s^-1^, 12 h photoperiod) at 28°C: 18°C (day: night) and 60 % relative humidity in a growth chamber (KuLan, China).

Tomato seedlings at the five-leaf stage were transferred to a dark growth chamber with an aerial temperature of 4 °C for 24 h (18:00-18:00 [the next day]) during LNT treatment, with 18 °C as the control temperature. All measurements were performed on the fourth fully expanded functional leaves during the experiment. Tomato leaves were collected to perform physiological experiments immediately after treatment, and the remaining leaves were collected in liquid nitrogen and stored at -80 °C for further physiological and molecular biology experiments. Five biologically independent replicates were collected for each treatment.

### Measurement of Pn, *F*_v_/*F*_m_, and NPQ

GFS-3000 and DUAL-PAM-100 synchronous measuring instruments (Heinz Walz, Effeltrich, Germany) were used to measure Pn in constant irradiation (228 μmol photons m^−2^ s^−1^, 635 nm). CO_2_ concentration was 600 ppm and the temperature approximately 25°C (Lu et al., 2020b). Tomato seedlings were first adapted to the dark for 30 min. *F*_v_/*F*_m_ and NPQ were measured using MAXI-Imaging-PAM (blue LED version) and the imaging fluorometer software Win (v2.46i, Heinz Walz, Effeltrich, Germany) at room temperature (approximately 25 °C). After initiating measurement, a saturating pulse (10,000 µmol photons m^−2^ s^−1^, 300 ms) was applied to obtain maximal fluorescence (*F*_m_). Both dark-adapted and light-adapted maximal fluorescence (*F*_m_ and *F*_m_’) were obtained with a saturating pulse under actinic light (228 µmol photons m^−2^ s^−1^, 635 nm). The weak modulated illumination light (<0.1 μmol m^-2^ s^-1^, frequency 0.6 kHz) was used to measure the dark-adapted and light-adapted initial fluorescence (*F*_o_ and *F*_o_’). The calculation formula is *F*_v_/*F*_m_ = (*F*_m_ - *F*_o_)/ *F*_m_, NPQ = (*F*_m_ - *F*_m_’)/ *F*_m_’.

### Analysis of H_2_O_2_, O_2_^−^, and ROS fluorescence

Accumulation of O_2_^−^ and H_2_O_2_ was detected using NBT and DAB staining. Fresh tomato leaves were dipped into a solution comprising 10 µg·mL^−1^ DAB and 0.2% NBT, respectively, incubated at room temperature for 12 h. Scanned and observed after decolorization.

Reactive oxygen species (ROS) fluorescence was observed under a confocal microscope (LSM880NLO, Zeiss, Germany) using 2’,7’-dichlorodihydro fluorescein diacetate (H2DCFDA, 10 InvM, Invitrogen, USA) as described by Yu et al. (2022). The excitation and absorption wavelengths of the ROS fluorescence were 488 nm and 525 nm, respectively.

### Total RNA extraction and qRT-PCR

Total RNA was extracted from tomato leaves using the plant total RNA extraction kit (Tiangen, Beijing, China) following the manufacturer’s instructions. First-strand complementary DNA (cDNA) was synthesized using a PrimeScript RT reagent kit with gDNA Eraser (TaKaRa, Dalian, China). SYBR Premix ExTaqTM (TaKaRa, Dalian, China), ABI 7500 Real-Time PCR system (Applied Biosystems, Foster City, CA), and Software 7500 v.2.0.6 were used for qRT-PCR and gene expression analysis. The primer sequences are listed in Table S1.

### Subcellular localization

The coding regions (without stop codons) of *SlPsbS*, *SlCSN5A,* and *SlCV* were within the BamHI and SalI sites upstream of GFP in the pCAMBIA1300 vector. *SlPsbS*, *SlCSN5A*, and *AtHPR2* (cytosol localized protein) (Ye et al., 2014) were ligated into the pCAMBIA1300-mCherry expression vector. The resulting construct was then transformed into *A. tumefaciens* strain EHA105 and infiltrated into the leaves of 3-week-old *N. benthamiana* plants. *N. benthamiana* plants were kept in the dark for 48 h after infiltration. Taking room temperature as the control, a few plants were treated with LNT for 24 h. The fluorescence signal was monitored with a confocal laser scanning microscope (LSM880NLO, Zeiss, Germany). For green fluorescence observation, the excitation wavelength was 488 nm and the emission wavelengths 520-540 nm. For red fluorescence observation, the excitation wavelength was 561 nm and the emission wavelengths 610-630 nm.

### Y2H assay

The SlCSN5A and SlCV coding sequences were inserted into the pGBKT7 vector, respectively, and the *SlPsbS* and *SlCSN5A* gene fragments into the pGADT7 vector, respectively. The fusion plasmid was co-transformed into the strain Y2H Gold. After transformation, yeast cells were grown on the SD-Trp-Leu (DDO) medium and transformants were grown on an SD-Trp-Leu-His-Ade (QDO) medium containing X-Gal (5-bromo-4-chloro-3-indolyl-b-D-galactopyranoside). Both 40 µg ml^-1^ of X-a-gal and 100 ng ml^-1^ aureobasidin A (AbA) were present in the QDO medium.

### BIFC assay

CDSs of *SlCSN5A* and *SlCV* were ligated into the pSPYNE-35S vector. CDSs of *SlPsbS* and *SlCSN5A* were amplified and cloned into the pSPYCE-35S vector. The resulting plasmids were introduced into *A. tumefaciens* strain EHA105, and then co-transformed into *N. benthamiana* leaves. YFP fluorescence was observed under a confocal laser scanning microscope (LSM880NLO, Zeiss, Germany) after 72 h. The excitation wavelength was 488 nm, and the emission wavelength 520-540 nm.

### Luciferase complementation imaging assays

CDSs of *SlPsbS* and *SlCSN5A* were inserted into the pCAMBIA1300-CLuc vector using the BamHI and PstI restriction sites. *A. tumefaciens* infiltration was carried out as above. LUC activity was detected 72 h after infiltration using the Night SHADELB 985 imaging system (Berthold Technologies, Germany). Thirty minutes before detection, 0.2 mM potassium luciferin (Gold Biotechnology Inc., St. Louis, MO, USA) was infiltrated into the same positions where *A. tumefaciens* was infiltrated.

### Immunoblot analysis

Plant leaf tissues were frozen in liquid nitrogen, and homogenized in an extraction buffer (Shang et al., 2019). Protein concentration was measured using a BCA Protein Assay Kit (Thermo Scientific, Waltham, MA, USA). Total proteins were loaded and separated on a 10% SDS-PAGE gel. Proteins were then transferred onto nitrocellulose membranes (BioRad, USA), which were incubated with anti-PsbS (PhytoAB, USA), anti-CSN5 (ABclonal, China) or anti-CV (prepared in this laboratory). After incubation with goat anti-rabbit or anti-mouse horseradish peroxidase (HRP)-linked antibodies (Solarbio, China), enhanced chemical luminescence (ECL) was performed to detect labeled proteins. Mouse anti-Actin polyclonal antibody (CWBIO, China) was used as a loading control.

Chloroplast proteins were extracted from tomato leaves using a commercial Chloroplast Protein Isolation Kit (BestBio, China) according to the manufacturer^’^s instructions. After separating the extracted chloroplast proteins using 10% SDS-PAGE gel, we conducted immunoblot analysis with anti-PsbS (PhytoAB, USA), anti-Actin (CWBIO, China), and anti-Ub (Proteintech, China) antibodies.

### Immunoprecipitation, protein degradation and ubiquitination assay

PCR-amplified fragments of *SlCSN5A* and *SlCV* CDSs were inserted into the plant expression vector pCAMBIA1300-GFP. CDSs of *SlPsbS* and *SlCV* were ligated into the pRI101-3хFLAG vector. *A. tumefaciens* strains carrying recombinant plasmid and p19 genes, as well as internal control plasmids, were infiltrated into *N. benthamiana* leaves either together or separately. Five days after infiltration, samples were collected for immunoblot analysis. Afterward, 50 μΜ cycloheximide (CHX, protein synthesis inhibitor in eukaryotes), 50 μΜ MG132 (26S proteasome inhibitor) or an equal concentration of DMSO were infiltrated before the samples were harvested according to the experimental needs (Liu et al., 2010).

*N. benthamiana* leaves co-expressing SlCSN5A-GFP/SlCV-GFP and SlPsbS-FLAG were collected for immunoprecipitation analysis. Total proteins were extracted in an IP buffer (Solarbio, China). The cell lysates were then incubated with anti-FLAG beads (AlpaLife, China) overnight at 4 °C in a top to end rotator. The samples were separated by SDS-PAGE and analyzed by immunoblot using anti-GFP (Solarbio, China) or anti-FLAG (Solarbio, China) antibodies. Further, the IP products of *N. benthamiana* leaves expressing SlPsbS-FLAG and SlCV-FLAG were immunodetected using an anti-Ub antibody (Proteintech, China) (Belda-Palazon et al., 2016; Luo et al., 2021).

### In vitro ubiquitination assay

Full-length sequences of *SlCSN5A* were inserted into the pGEX-4T-1 vector. The resulting plasmids were transformed into *Escherichia coli* BL21 (DE3) competent cells. The recombinant protein was then purified by GST-tag Purification Resin (Beyotime, China). The purified product was co-incubated with E1 (ubiquitin-activating enzyme), E2 (ubiquitin-conjugating enzyme), and ubiquitin (Ub) (30 °C, 1.5 h). They were then separated on an SDS-PAGE gel, transferred to membranes, and immunoblotted against anti-GST antibodies (Solarbio, China). Anti-Ub antibody (Proteintech, China) was used to detect ubiquitin.

### Pull-down assay

The CDSs of SlCSN5A, SlPsbS, and SlCV were separately ligated into pGEX4T-1 and pET-32a for the expression of fusion proteins in E. coli. Mix the appropriate amount of recombinant protein in binding buffer (50 mM Tris-HCl, PH=7.5, 100 mM NaCl, 0.25% (v/v) Triton X-100, 35 mM β-mercaptoethanol) and incubate at 4°C for 4h. Add 50 μl of GST-tag Purification Resin (Beyotime, China) for 2h at 4°C with rotation. Subsequently, nonspecifically bound proteins were removed by washing 5 times with buffer. The target protein was eluted and detected via immunoblot analysis. GST protein was used as the negative control.

### TUBE assay

The seedlings were collected and ground in liquid nitrogen. The resulting powders were then mixed with extraction buffer (50 mM Tris-HCl, pH 7.5, 150 mM NaCl, 0.1% (v/v) NP-40, 1% (v/v) Triton X-100,1 mM PMSF, 1х complete protease inhibitor cocktail, 5 mM N-ethylmaleimide, and 50 µM MG132), and the cellular debris was removed by two rounds of centrifugation at 20,000 g at 4 °C. TUBE2 proteins were added into the supernatant and incubated in the dark at 4 °C overnight. Then 50 µL glutathione agarose beads were added and incubated at 4 °C for 1 h. The captured fractions were then washed with wash buffer (50 mM Tris-HCl, pH 7.5, 500 mM NaCl, 0.1% (v/v) NP-40, 1% (v/v) Triton X-100) three times. Western blot analysis using anti-PsbS and anti-Ub antibodies.

### Statistical analysis

A completely randomized design (CRD) was used for the experiments. Data were analyzed using the SPSS v.22 software (IBM SPSS STATISTICS, USA). A value of P <0.05 was considered to indicate statistical significance. Figures were drawn with the Origin 2022 software (Origin Lab, Northampton, MA, USA). Primers used for vector constructs are listed in Table S2.

## ACKNOWLEDGEMENTS

We are grateful to Taotao Wang (Huazhong Agricultural University) for providing the *SlCSN5*-RNAi plants, Yangwen Qian (Hangzhou Biogle Co.Ltd) for tomato transformation, Mingnan Qu (Hainan Yazhou Bay Seed Laboratory) and Jialong Li (Institute of Botany, Chinese Academy of Sciences) for the experimental method. The authors would like to thank TopEdit (www.topeditsci.com) for its linguistic assistance during the preparation of this manuscript. This work was supported by the National Natural Science Foundation of China (Grant No. 32272791; 32072651), the earmarked fund for CARS (CARS-23), Joint Fund for Innovation Enhancement of Liaoning Province (2021-NLTS-11-01), and the support program for Young and middle-aged Scientific and Technological Innovation Talents (RC210293).

## AUTHOR CONTRIBUTIONS

Y.L. and T.L. conceived and designed the project. J.L., J.Y., P.L., J.G., Y.C., T.Z., and J.L. performed the experiments. T.W. created the tomato *SlCSN5*-RNAi materials. J.L., and Y.L. wrote the manuscript. W.Y., R.L., F.W., and M.Q. revised and edited the manuscript.

## Notes

### Competing Interest Statement

The authors have declared no competing interest.

