## Supplemental information for "Ubiquitin-mediated stabilization of SlPsbS regulates low night temperature tolerance in tomatoes"

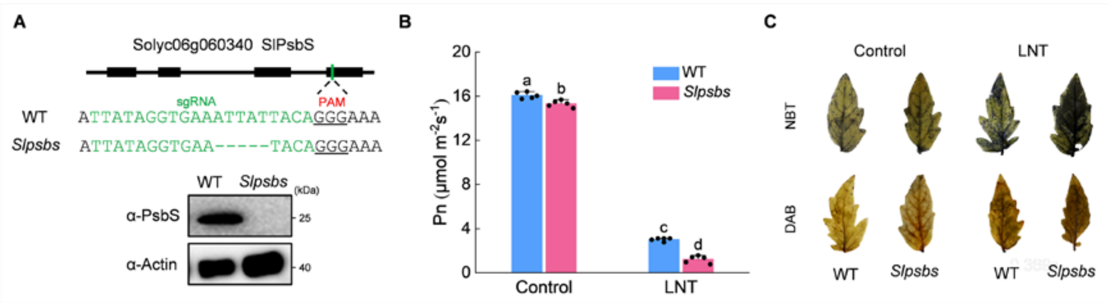


**Figure S1.** SlPsbS silencing increases the sensitivity of tomato plants to LNT stress. (A) *Slpsbs* mutant identification results and PsbS protein levels analysis in WT and *Slpsbs* mutants. Immunoblotting with anti-Actin antibody provided control for equal loading. The numbers on the right denoted the molecular mass of marker proteins in kiloDaltons. (B) Net photosynthetic rate (Pn). Data are the means of five replicates with standard errors shown by vertical bars. Differences among treatments were analyzed by the one-way ANOVA comparison test ($P < 0.05$). Different letters indicate significant differences among treatments. (C) Histochemical staining of tomato leaves with tetranitroblue tetrazolium chloride (NBT) and diaminobenzidine (DAB).

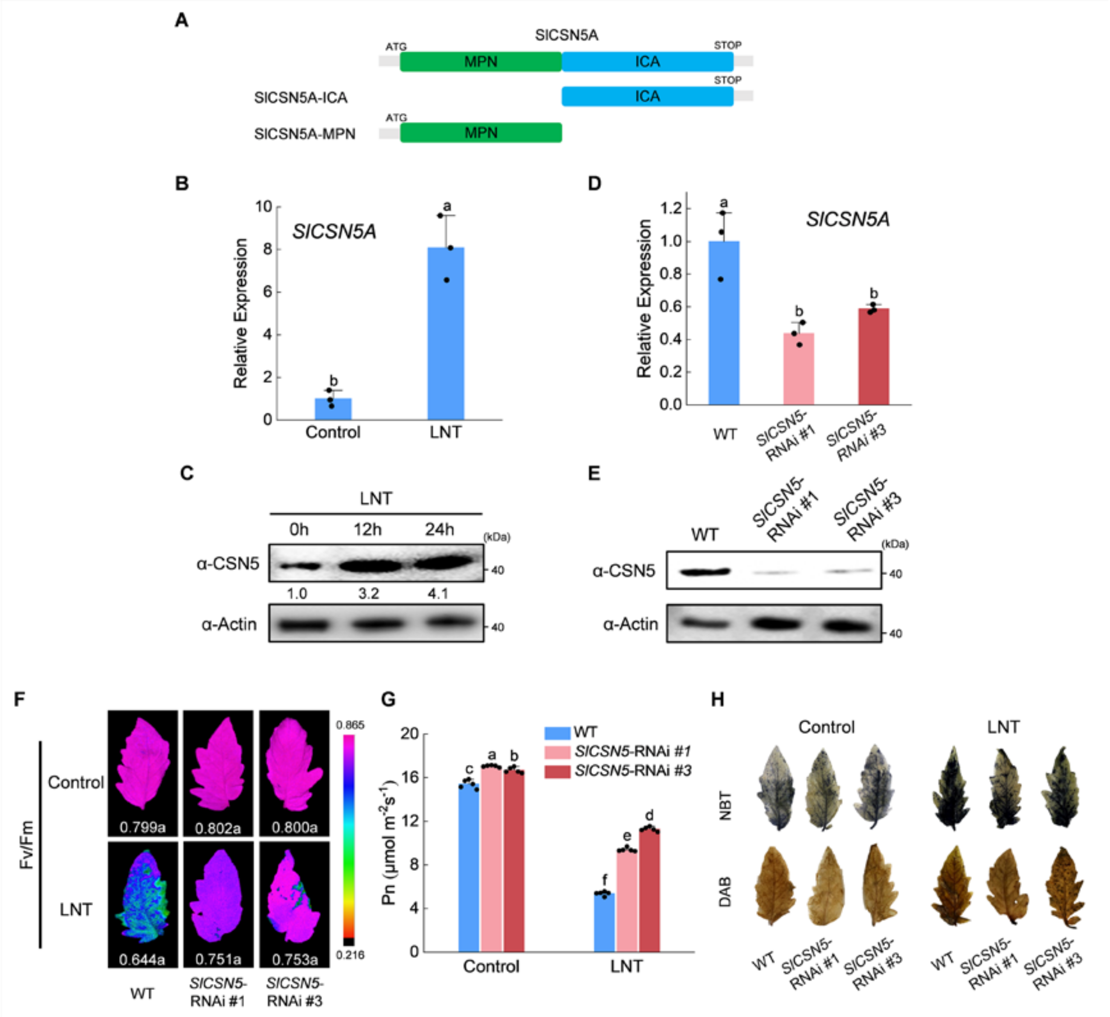


**Figure S2.** (A) Diagram of SlCSN5A gene structure. (B) Relative expression of *SlCSN5A* under LNT stress. (C) The protein level of SlCSN5 under LNT stress. (D)

SlCSN5 protein levels in WT and *SlCSN5*-RNAi line plants. (E) The protein level of SlCSN5 in WT and *SlCSN5*-RNAi line plants. Immunoblotting with anti-Actin antibody provided control for equal loading. Intensities of bands were quantified by ImageJ normalized to Actin and expressed relative to controls. The relative amounts are shown below each lane. The numbers on the right denoted the molecular mass of marker proteins in kiloDaltons. (F) The maximum quantum yield of PSII ($F_v/F_m$). (G) Net photosynthetic rate (Pn). Data are the means of five replicates with standard errors shown by vertical bars. Differences among treatments were analyzed by the one-way ANOVA comparison test ($P < 0.05$). Different letters indicate significant differences among treatments. (H) Histochemical staining of tomato leaves with tetranitroblue tetrazolium chloride (NBT) and diaminobenzidine (DAB).

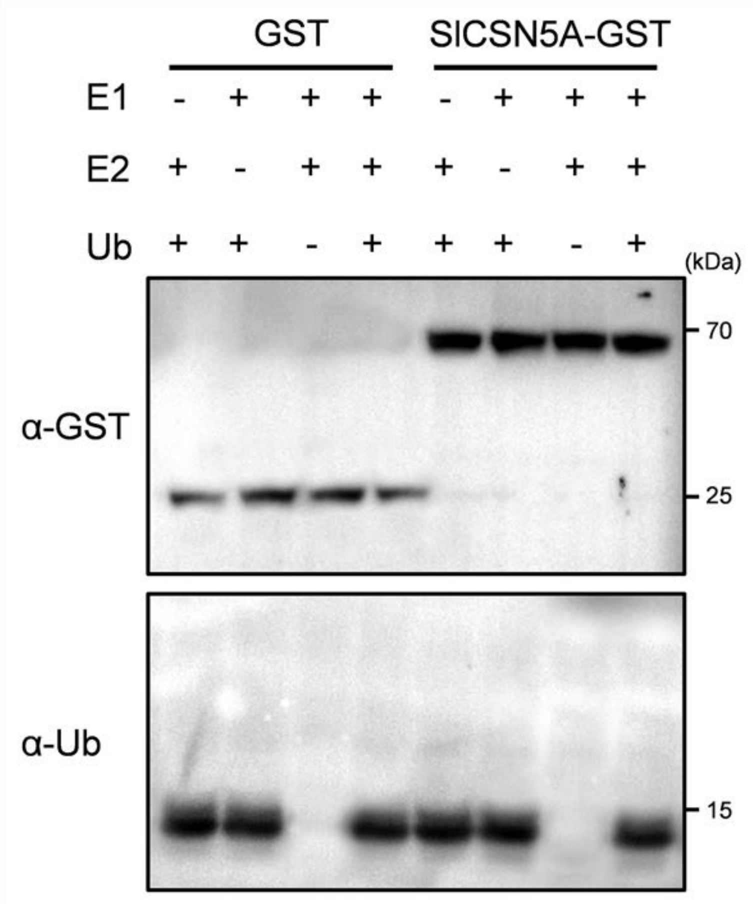


**Figure S3.** SlCSN5A does not have E3 ubiquitin ligase activity. Ubiquitination reactions in vitro. The SlCSN5A-GST fusion protein was assayed for ubiquitin activity in the presence or not presence of E1, E2, and/or ubiquitin. The immunoblotted against anti-GST antibody. The anti-Ub antibody was used to detect ubiquitin. E1, ubiquitin-activating enzyme. E2, ubiquitin-conjugating enzyme. Ub, ubiquitin. The numbers on the right denoted the molecular mass of marker proteins in kiloDaltons.

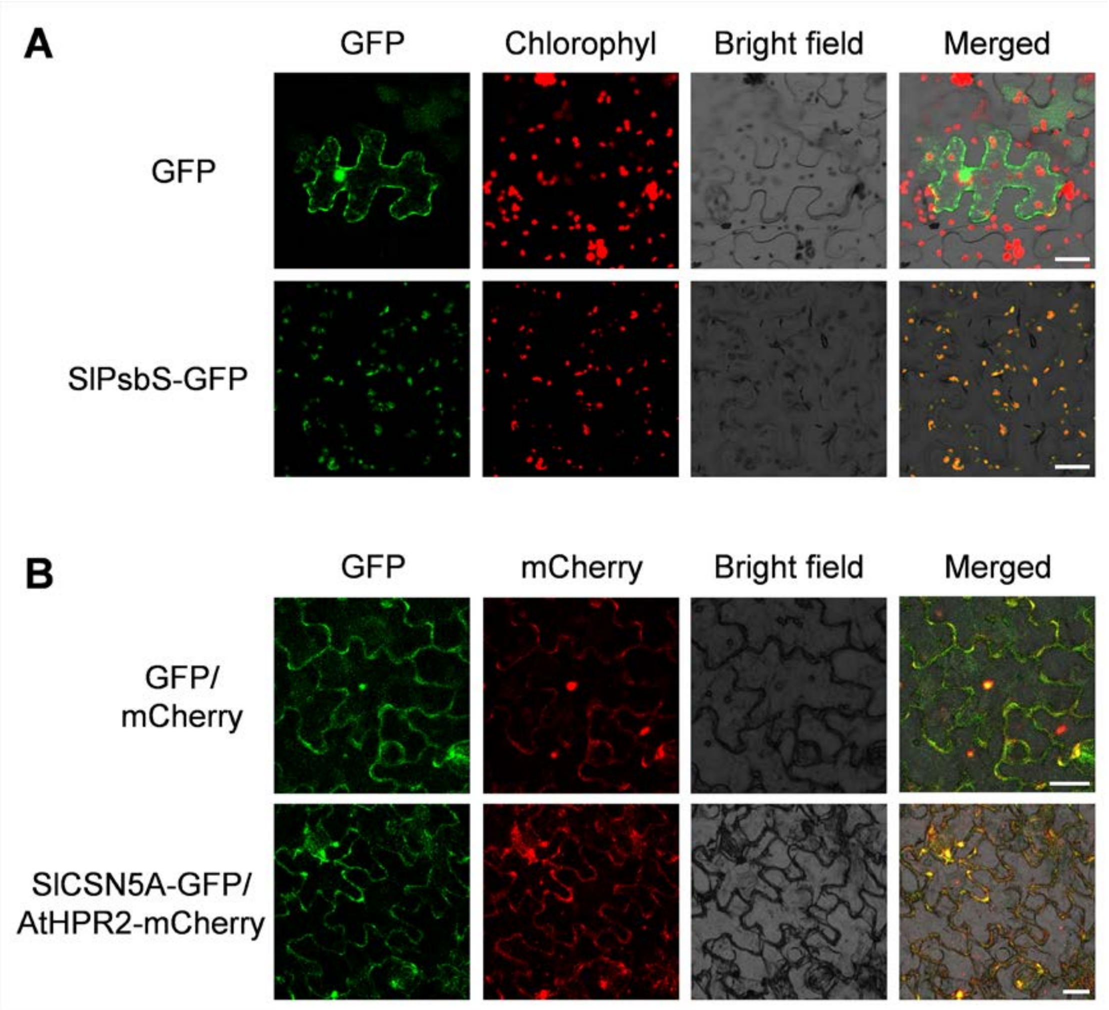


**Figure S4.** Subcellular localization of SlPsbS and SlCSN5A. (A) SlPsbS and (B) SlCSN5A. HPR2-mCherry was used as a cytosol marker (bar, 25 μm).

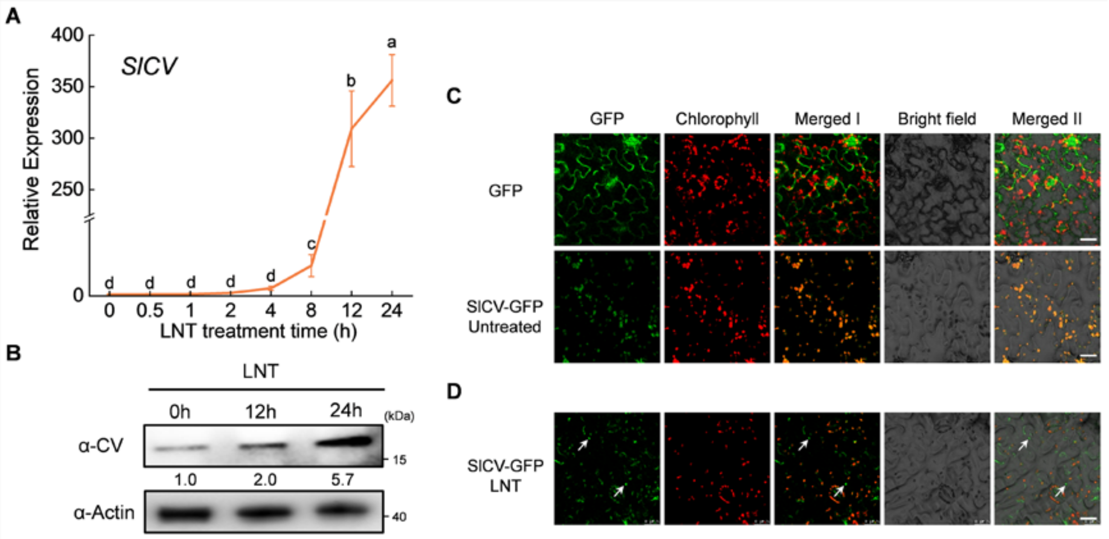


**Figure S5.** (A) Relative expression of *SlCV* under LNT stress. (B) The protein level of SlCV under LNT stress. Immunoblotting with anti-Actin antibody provided control for equal loading. Intensities of bands were quantified by ImageJ and normalized to Actin and expressed relative to controls. The relative amounts are shown below each lane. The numbers on the right denoted the molecular mass of marker proteins in kiloDaltons. (C) Subcellular localization of SlCV. *A. tumefaciens* strain EHA105 containing the SlCV-GFP construct was infiltrated into *N. benthamiana* leaves and examined after 72 h. (D) SlCV-GFP construct infiltrated *N. benthamiana* leaves for 48h and then treated with LNT for 24h to observe the fluorescence signal (bar, 25 μm).

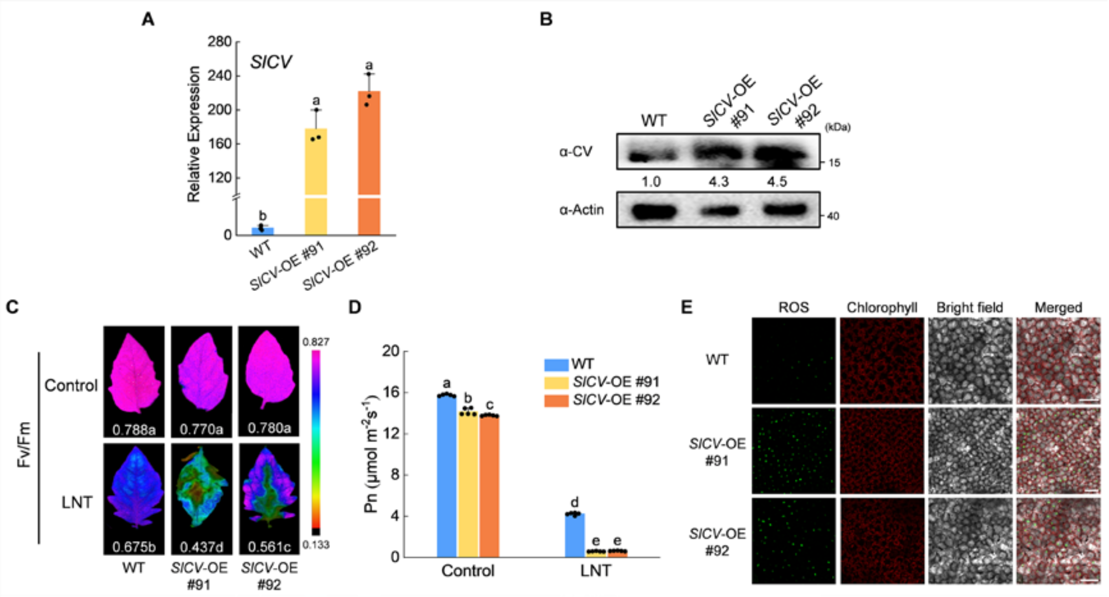


**Figure S6.** (A) Relative expression of *SlCV* in WT and *SlCV* overexpression plants. (B) SlCV protein levels were analyzed by immunoblot analysis in WT and *SlCV* overexpression plants. Immunoblotting with anti-Actin antibody provided control for equal loading. Intensities of bands were quantified by ImageJ and normalized to Actin and expressed relative to controls. The relative amounts are shown below each lane. The numbers on the right denoted the molecular mass of marker proteins in kiloDaltons. (C) The maximum quantum yield of PSII ($F_v/F_m$). (D) Net photosynthetic rate (Pn). Data are the means of five replicates with standard errors shown by vertical bars. Differences among treatments were analyzed by the one-way ANOVA comparison test ($P < 0.05$). Different letters indicate significant differences among treatments. (E) ROS fluorescence (stained with H2DCFDA, ROS) in the *SlCV* overexpression and WT plants under LNT stress (bar, 100 μm).

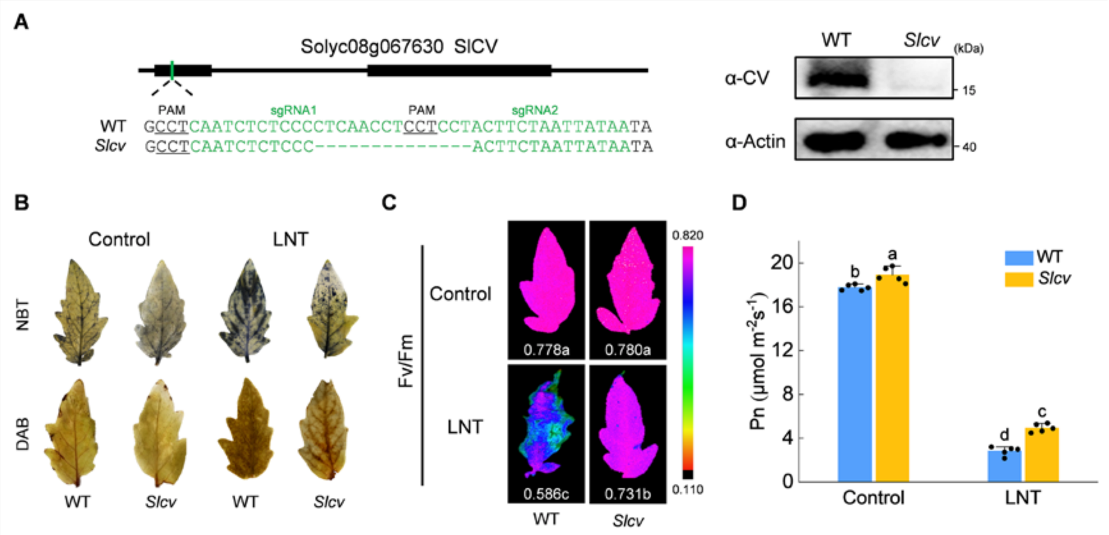


**Figure S7.** (A) *Slcv* mutant identification result and CV protein levels analysis in WT and *Slcv* mutants. Immunoblotting with anti-Actin antibody provided control for equal loading. The numbers on the right denoted the molecular mass of marker proteins in kiloDaltons. (B) Histochemical staining of tomato leaves with tetranitroblue tetrazolium chloride (NBT) and diaminobenzidine (DAB). (C) The maximum quantum yield of PSII ($F_v/F_m$). (D) Net photosynthetic rate (Pn). Data are the means of five replicates with standard errors shown by vertical bars. Differences among treatments were analyzed by the one-way ANOVA comparison test (P < 0.05). Different letters indicate significant differences among treatments.

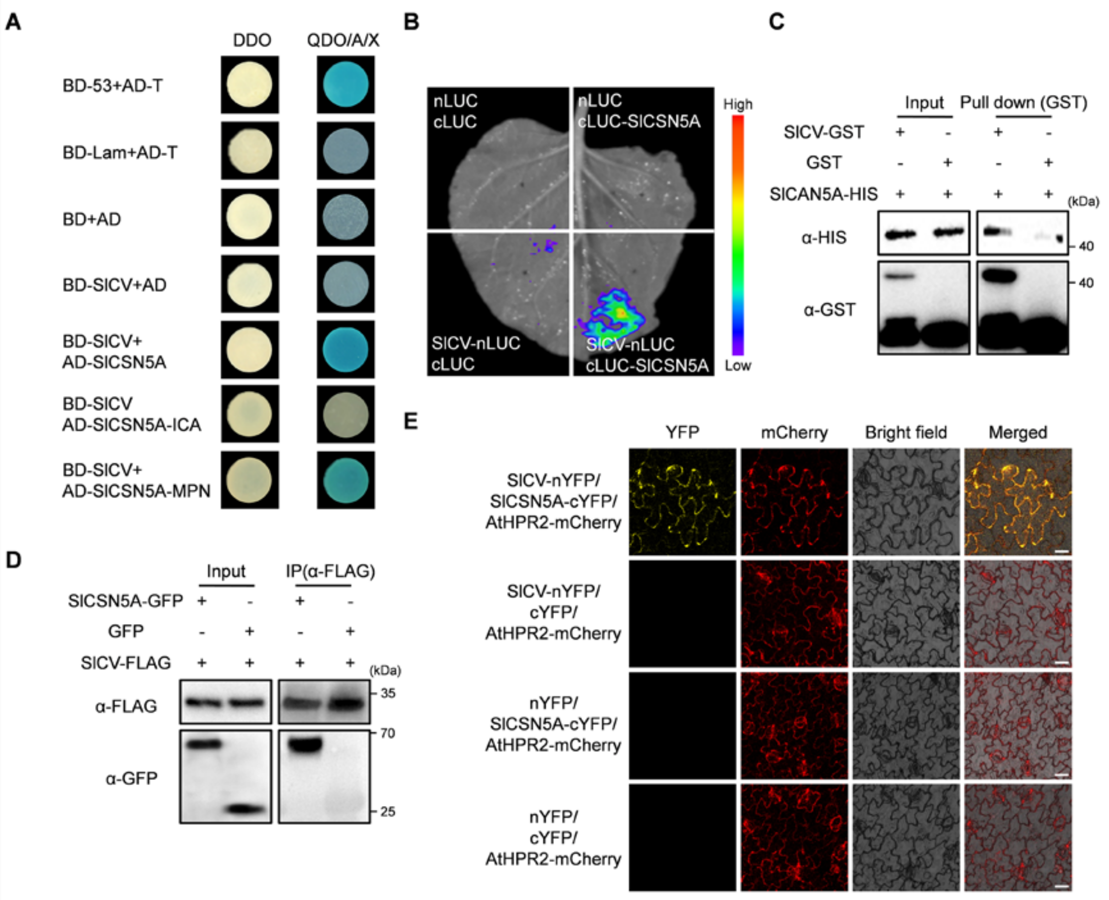

**Figure S8.** SlCSN5A interact with SlCV. (A) Yeast two-hybrid (Y2H) assays showing the interaction between SlCV and SlCSN5A. SlCV and SlCSN5A were fused to either the DNA binding domain (BD) or the activation domain (AD) of GAL4. DDO, SD medium lacking Trp/Leu; QDO/A/X, SD medium lacking Trp/Leu/His/Ade and containing X-α-gal and aureobasidin A. The empty plasmids were used as controls. The blue color indicates protein interaction. (B) Luciferase complementation imaging assay showing that SlCV interaction with SlCSN5A in *N. benthamiana* leaves. *A. tumefaciens* strain EHA105 harboring different constructs was infiltrated into *N. benthamiana* leaves and examined after 72 h. (C) Pull-down assay showing that SlCV interacts with SlCSN5A in vitro. The SlCV-GST and SlCSN5A-HIS fusion proteins were expressed in *E. coli* and purified. SlCV-GST was bound to GST magnetic beads. The assays were analyzed using immunoblotting with anti-GST and anti-HIS antibodies. (D) Co-immunoprecipitation assay. SlCSN5A-GFP and SlCV-FLAG were cotransformed in *N. benthamiana* leaves. The leaves were collected and extracted total protein, and immunoprecipitation against anti-FLAG beads. Immunoblot analysis with anti-GFP and anti-FLAG antibodies. The numbers on the right denoted the molecular mass of marker proteins in kiloDaltons. (E) Interaction of SlCV and SlCSN5A detected by bimolecular fluorescence complementation (BIFC) analysis. SlCV-nYFP and SlCSN5A-cYFP were co-infiltrated into *N. benthamiana* leaves and expressed for 72 h. HPR2-mCherry was used as a cytosol marker (bar, 25 μm).

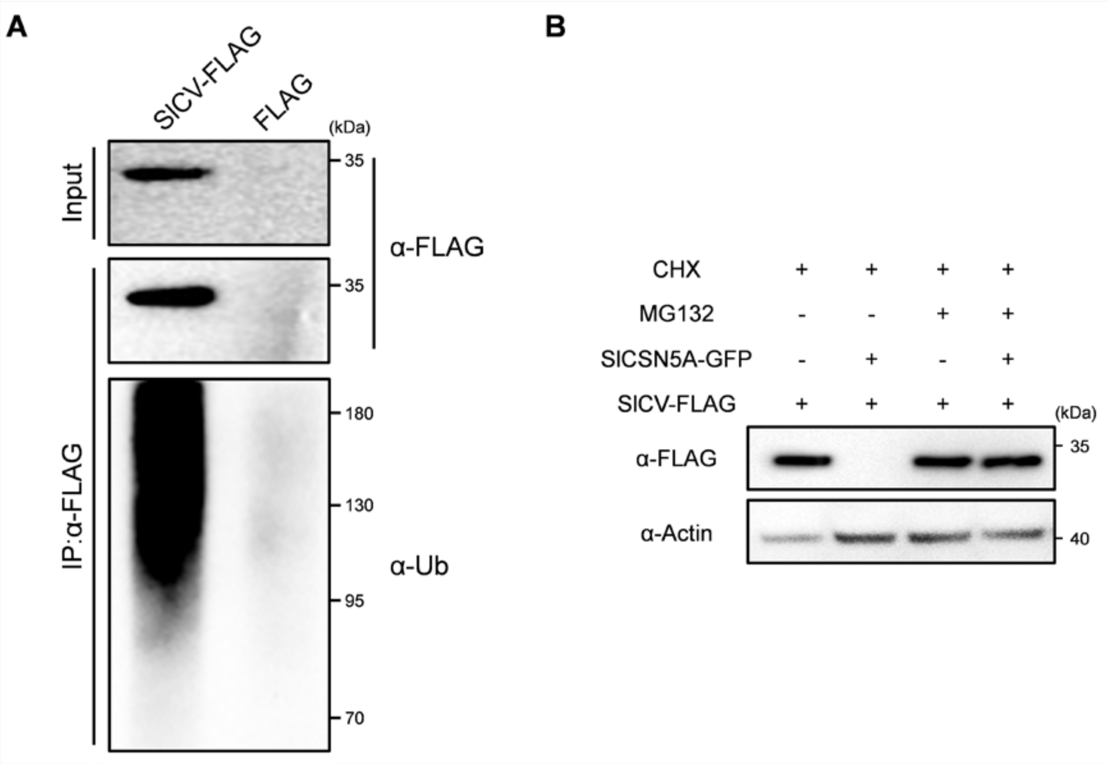


**Figure S9.** SlCV can be ubiquitinated. (A) The SlCV-FALG plasmid was transiently into *N. benthamiana* leaves and immunoprecipitated with anti-FLAG beads. Immunoblot analysis was performed using anti-FLAG and anti-Ub antibodies. (B) SlCSN5A promotes SlCV degradation by the 26s proteasome. The SlCSN5A-GFP and SlCV-FLAG construct combinations were cotransformed into *N. benthamiana* leaves. Five days after infiltration immunoblotted against anti-FLAG antibody. CHX, protein synthesis inhibitor in eukaryotes. MG132, 26S proteasome inhibitor. The anti-Actin served as a loading control and numbers on the right denoted the molecular mass of marker proteins in kiloDaltons.

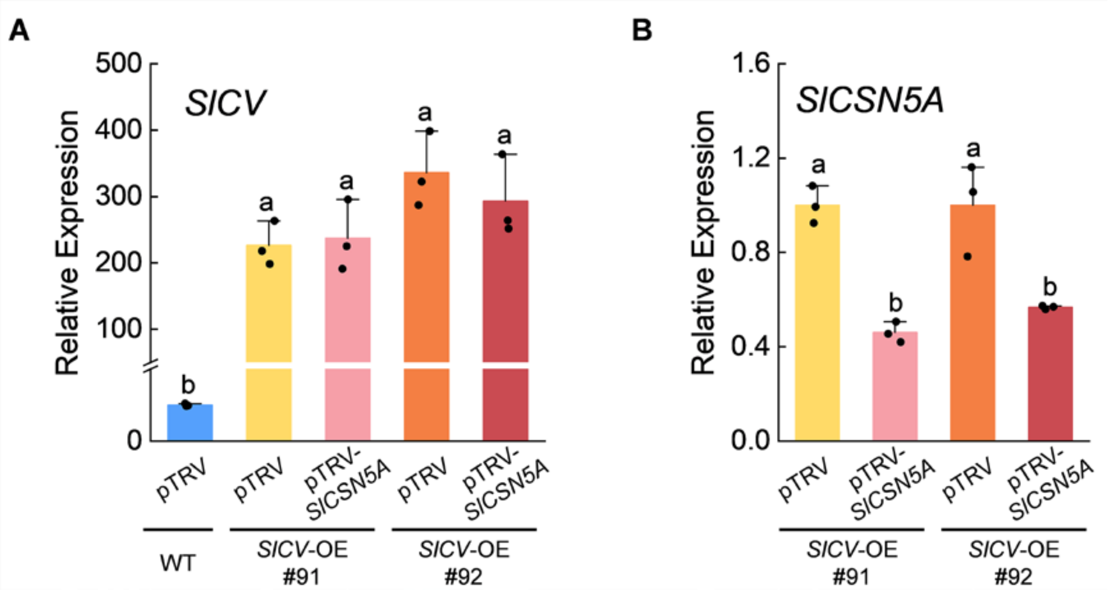


**Figure S10.** Relative expression of (A) *SlCV* and (B) *SlCSN5A* of silenced (pTRV-*SlCSN5A*) in *SlCV* -OE plants and nonsilenced *SlCSN5A* (pTRV) in WT and CV-OE plants. Data are the means of five replicates with standard errors shown by vertical bars. Differences among treatments were analyzed by the one-way ANOVA comparison test (P < 0.05). Different letters indicate significant differences among treatments.

**Table S1 List of primer sequences used for qRT-PCR analysis.**

| Gene | Accession number | Primer sequences (5'-3') |
|---|---|---|
| *SlCSN5A* | Solyc06g073150 | F-CAGCAAACCTGGGAGTTAGAGA |
| | | R-TGAGAAGAGCAAGAGCAGAGATC |
| *SlCV* | Solyc08g067630 | F-TCCCCTCAACCTCCTCCTACT |
| | | R-TGGGCATGATCTTTTTTCACTC |
| *ACTIN* | Solyc11g005330 | F-TGTCCCTATTTACGAGGGTTATGC |
| | | R-CAGTTAAATCACGACCAGCAAGAT |

**Table S2 PCR primer sequences used for vector construction.**

| Vector | Primer sequences (5’-3’) |
|---|---|
| SlPsbS-GFP | F-GAGCTCGGTACCCGGGGATCC ATGGCTCAAACAATGTTGTT<br>R-GCCCTTGCTCACCATGTCGAC ATCTTCTTCCTCATCAGTGAT |
| SlCSN5A-GFP | F-GAGCTCGGTACCCGGGGATCC ATGGACTCTCTGAATTCTTACGCATCGT<br>R-GCCCTTGCTCACCATGTCGAC GCTTTCGATCATGGGCTCGG |
| SlCV-GFP | F-GAGCTCGGTACCCGGGGATCC ATGGCTATTTCAACAAAGTTCTGC<br>R-GCCCTTGCTCACCATGTCGAC CATAGTGAAACATCCTTTACTAAAT |
| SlCSN5A-mCherry | F-GCTTCGAATTCTGCAGTCGAC ATGGACTCTCTGAATTCTTACGCATCGT<br>R-GCCCTTGCTCACCATCAGGAT GCTTTCGATCATGGGCTCGG |
| SlPsbS-mCherry | F-GCTTCGAATTCTGCAGTCGAC ATGGCTCAAACAATGTTGTT<br>R-GCCCTTGCTCACCATCAGGAT ATCTTCTTCCTCATCAGTGAT |
| SlCSN5A-BK | F-ATGGCCATGGAGGCCGAATTC ATGGACTCTCTGAATTCTTACGCAT<br>R-TAGTTATGCGGCCGCTGCAGG TCAGCTTTCGATCATGGGCT |
| SlCSN5A-AD | F-GCCATGGAGGCCAGTGAATTC ATGGACTCTCTGAATTCTTACGCAT<br>R-AGCTCGAGCTCGATGGATCCC TCAGCTTTCGATCATGGGCT |
| SlPsbS-AD | F-GCCATGGAGGCCAGTGAATTC ATGGCTCAAACAATGTTGTTAAC<br>R-AGCTCGAGCTCGATGGATCCC CTAATCTTCTTCCTCATCAGTGATA |
| SlCV-BK | F-ATGGCCATGGAGGCCGAATTC ATGGCTATTTCAACAAAGTTCTGC<br>R-TAGTTATGCGGCCGCTGCAGG CATAGTGAAACATCCTTTACTAAAT |
| SlCSN5A-MPN-AD | F-GCCATGGAGGCCAGTGAATTC ATGGACTCTCTGAATTCTTAC<br>R-CAGCTCGAGCTCGATGGATCC AAGAAAGGGTTCCTGATATT |
| SlCSN5A-MPN-BK | F-ATGGCCATGGAGGCCGAATTC ATGGACTCTCTGAATTCTTAC<br>R-TAGTTATGCGGCCGCTGCAG AAGAAAGGGTTCCTGATATT |
| SlCSN5A-ICA-AD | F-GCCATGGAGGCCAGTGAATTC GCAGTTGTTATTGATCCAACAAGAAC<br>R-CAGCTCGAGCTCGATGGATCC TCAGCTTTCGATCATGGGCT |
| SlCSN5A-ICA-BK | F-ATGGCCATGGAGGCCGAATTC GCAGTTGTTATTGATCCAACAAGAAC<br>R-TAGTTATGCGGCCGCTGCAG TCAGCTTTCGATCATGGGCT |
| SlCSN5A-NE | F-CCCAGGCCTACTAGTGGATCC ATGGACTCTCTGAATTCTTACGCAT<br>R-GGGAAATTCGAGCTCCTACCC TCAGCTTTCGATCATGGGCT |
| SlCV-NE | F-CCCAGGCCTACTAGTGGATCC ATGGCTATTTCAACAAAGTTCTGC<br>R-GGGAAATTCGAGCTCCTACCC CATAGTGAAACATCCTTTACTAAAT |
| SlCSN5A-CE | F-TGGCGCGCCACTAGTGGATCC ATGGACTCTCTGAATTCTTACGCAT<br>R-AACATCGTATGGGTACATCCC GCTTTCGATCATGGGCTCG |
| SlPsbS-CE | F-TGGCGCGCCACTAGTGGATCC ATGGCTCAAACAATGTTGTTAAC<br>R-AACATCGTATGGGTACATCCC CTAATCTTCTTCCTCATCAGTGATA |
| | F-CCGGGGCGGTACCCGGGAT ATGGACTCTCTGAATTCTTACGCAT |

| | |
|---|---|
| SlCSN5A-CLUC | R-ATACGAACGAAAGCTCTGCAG TCAGCTTTCGATCATGGGCT |
| SlCSN5A-NLUC | F-CGAGCTCGGTACCCGGGATCC ATGGACTCTCTGAATTCTTACGCAT<br>R-CGCGTACGAGATCTGGTCGAC GCTTTCGATCATGGGCTCG |
| SlCV-NLUC | F-CGAGCTCGGTACCCGGGATCC ATGGCTATTTCAACAAAGTTCTGC<br>R-CGCGTACGAGATCTGGTCGAC CATAGTGAAACATCCTTTACTAAAT |
| SlPsbS-FLAG | F-TCTTCACTGTTGATACATATG ATGGCTCAAACAATGTTGTTAAC<br>R-AGAGTTGTTGATTCAGAATTCTTAAGCGTAATATGGAACATCGT CTAATCTTCTTCCTCATCAGTGATA |
| SlCV-FLAG | F-TCTTCACTGTTGATACATATG ATGGCTATTTCAACAAAGTTCTGC<br>R-AGAGTTGTTGATTCAGAATTCTTAAGCGTAATATGGAACATCGT ATGGGTACAT CATAGTGAAACATCCTTTACTAAAT |
| SlCSN5A-GST | F-TTCCAGGGGCCCCTGGGATCC ATGGACTCTCTGAATTCTTACGCAT<br>R-GTCACGATGCGGCCGCTCGAG TCAGCTTTCGATCATGGGCT |
